# Division and adaptation to host nutritional environment of apicomplexan parasites depend on apicoplast lipid metabolic plasticity and host organelles remodelling

**DOI:** 10.1101/585737

**Authors:** Souad Amiar, Nicholas J. Katris, Laurence Berry, Sheena Dass, Melanie J. Shears, Camille Brunet, Bastien Touquet, Mohamed-Ali Hakimi, Geoffrey I. McFadden, Yoshiki Yamaryo-Botté, Cyrille Y. Botté

## Abstract

Apicomplexan parasites are unicellular eukaryotes responsible for major human diseases including malaria and toxoplasmosis. Apicomplexan parasites must obtain and combine lipids both from host cell scavenging and *de novo* synthesis to maintain parasite propagation and survival within their human host. Major questions on the actual role for each lipid source or how these are regulated upon fluctuating host nutritional conditions remain unanswered. Characterization of an apicoplast acyltransferase TgATS2, shows that the apicoplast provides local (lyso)phosphatidic acid balance, which is required for the recruitment of a novel dynamin (TgDrpC) critical during parasite cytokinesis. Disruption of TgATS2 led parasites to shift metabolic lipid acquisition from *de novo* synthesis towards host scavenging. Importantly, both lipid scavenging and *de novo* synthesis pathways exhibit major metabolic and cellular plasticity upon sensing host lipid-deprived environments through concomitant (i) up-regulation of *de novo* fatty acid synthesis capacities in the apicoplast, and (ii) parasite-driven host cell remodelling to generate multi-membrane-bound structures from host organelles that are imported towards the parasite.

## Introduction

Apicomplexa are protozoa, most of which are intracellular parasites that cause serious infectious and chronic diseases in humans including malaria, and toxoplasmosis. Most Apicomplexa harbour a relict non-photosynthetic plastid, the apicoplast, acquired by the secondary endosymbiosis of a red alga. The apicoplast lost photosynthetic capability during the conversion to a parasitic lifestyle (Botté et al., 2011; Botte et al., 2013). However, it still contains plant-like pathways including a prokaryotic type II fatty acid synthesis pathway (FASII) (Waller et al., 1998). The apicoplast is essential for parasite development and survival in both *T. gondii* and *P. falciparum* (MacRae et al., 2012).

The FASII pathway is thought to be essential for parasite survival only during specific life stages (Waller et al., 1998). Indeed in *Plasmodium*, disruption of FASII demonstrated that it is dispensable in asexual blood stages but essential for late liver stage in rodent malaria parasites, and for sporozoite schizogony during mosquito stages (Vaughan et al., 2009). Nevertheless, changes in *P. falciparum* blood stage growth conditions—such as lipid starvation during *in vitro* growth, or physiological stress in human patients—induced re-activation of apicoplast FASII (Botte et al., 2013; Daily et al., 2007), suggesting plasticity of the FASII in response to nutritional environment. In *T. gondii* FASII is essential during tachyzoite development, including for apicoplast biogenesis and parasite proliferation (Mazumdar et al., 2006; Ramakrishnan et al., 2015). Taken together, these data suggest that fatty acids (FA)/phospholipids (PL) produced by apicoplast FASII are a vital contribution to parasite membranes generated during intense stages of parasite replication and division in their life cycle.

Apicomplexan parasite membranes comprise up to 80% PL as a proportion of total lipid classes, primarily phosphatidylcholine (PC), phosphatidylethanolamine (PE), phosphatidylserine (PS) and phosphatidylinositol (PI, (Gulati et al., 2015; Welti et al., 2007). *T. gondii* can readily scavenge PL, and triacylglycerols (TAG) from the host but is also capable of-, and dependent on-*de novo* synthesis of several PL classes (Amiar et al., 2016; Charron and Sibley, 2002; Hu et al., 2017; Nolan et al., 2017). Like other eukaryotes, apicomplexan *de novo* PL synthesis is initiated by the assembly of FA (i.e. esterification onto a glycerol-phosphate backbone) into specific PL precursors. In *T. gondii*, FAs to be integrated into PLs derive from three not exclusive sources: (i) apicoplast FASII generating FA chains (C12:0, C14:0, and C16:0) (Amiar et al., 2016; Ramakrishnan et al., 2012), (ii) FA elongases located on the parasite endoplasmic reticulum (ER; C16:1, C18:1, C20:1, C22:1 and C26:1) (Dubois et al., 2018; Ramakrishnan et al., 2015), and (iii) FAs scavenged from the host cell (Bisanz et al., 2006; Charron and Sibley, 2002).

Typically, phosphatidic acid (PA) is the central precursor for the *de novo* synthesis of all phospholipid classes by the two-step esterification of FAs onto a glycerol-3-phosphate backbone; first by glycerol-3-phosphate acyltransferases (GPAT) to form lysophosphatidic acid (LPA), and then by acyl-glycerol-3-phopsphate acyltransferases (AGPAT) to convert LPA to PA. In eukaryotic cells, GPATs and AGPATs of diverse origins (prokaryotic, eukaryotic, plant/algal-like) work as a pair at several locations within the cell. Apicomplexan seem have two pairs of acyltransferases: one plastid-like pair putatively in the apicoplast, and another pair predicted to work in the ER (Amiar et al., 2016; Lindner et al., 2014; Shears et al., 2017). In *T. gondii* the apicoplast GPAT, *Tg*ATS1, is essential for tachyzoite development where it generates LPA from apicoplast-FA for the bulk synthesis of PL (Amiar et al., 2016). Recently, PA was shown to act as an essential signal transducer for microneme secretion in *T. gondii* (Bullen et al., 2016). Beyond their roles as lipid precursors and signal transducers, PA (and LPA) also have important biophysical properties by controlling the formation of positive or negative membrane curvatures, and thereby influence the recruitment of proteins involved in membrane fusion/fission events in other eukaryotic models (Brown et al., 2008; Kooijman et al., 2005; Schmidt et al., 1999).

Lipidomics suggests that most *T. gondii* PLs are hybrids, comprising one FA moiety from the apicoplast *de novo* synthesis pathway and a second one scavenged from the host (Amiar et al., 2016). Thus, both scavenging and *de novo* synthesis of FA are critical for intracellular development. Recent studies suggest that host nutritional changes can impact metabolism, intracellular development, and thus parasite virulence or stage differentiation, especially in *P. falciparum* (Brancucci et al., 2017; Costa et al., 2018; Mancio-Silva et al., 2017; Zuzarte-Luis et al., 2017). However, regulation of these pathways is poorly understood and nothing is known on the sensing of host environmental/nutritional changes in *T. gondii*.

Here we characterise *T. gondii* AGPATs, focusing on one (*Tg*ATS2) that we localize in the apicoplast. We show the function of *Tg*ATS2 as an acyltransferase by complementing a bacterial mutant, and we construct a knockdown mutant in which we observe the impact on parasite division, and the parasite lipid profile. Particularly the impact of PA/LPA changes on the localization of a dynamin-related protein (*Tg*DrpC) in the *Tg*ATS2 mutant is described and provides a rationale for cytokinesis defects associated with drug inhibition of apicoplast FASII (Martins-Duarte et al., 2015). Finally, changes in parasite lipid composition and lipid fluxes led us to subject parasites to lipid starvation to explore how host nutritional environment impacts parasite growth. Analysis of lipid fluxes and growth screening under adverse lipid conditions shows that parasites can sense the environment and respond by (i) upregulation of *de novo* lipid synthesis in the apicoplast and (ii) manipulate the host through vesiculation from host organelles and import of such material to the parasitophorous vacuole (PVM) mediated by export of parasite effectors to improve their lipid scavenging.

## Results

### Deletion of the *Toxoplasma gondii* apicoplast acyltransferase *Tg*ATS2 results in replication and division defects due to aberrant cytokinesis and residual body formation

To explore *de novo* PA synthesis in *T. gondii*, we searched the genome for AGPATs potentially able to catalyse the esterification of an activated FA (i.e acyl-CoA or acyl-ACP) onto LPA to make PA. We found two AGPAT candidates with conserved motifs: TGME49_297640, and TGME49_240860. Phylogenetic analyses reveal that TGME49_297640 clusters with a prokaryotic clade with plant and algal sequence and that TGME49_240860 clusters with a eukaryotic clade (**Fig. 1AB**). So we termed these enzymes *Tg*ATS2, and *Tg*AGPAT based on the plant and eukaryotic terminology, respectively. We generated parasite lines expressing *Tg*ATS2 and TgAGPAT endogenously tagged at the C-terminus with a triple HA tag, under control of their respective promoters (**Fig. S1**). Immunofluorescence assays (IFAs) confirmed that *Tg*ATS2 targets to the apicoplast (**Fig. 2A**), and that transient expression of TgAGAPT-HA showed a perinuclear structure that corresponds to the parasite ER (**Fig. 2B**).

**Figure 1:**
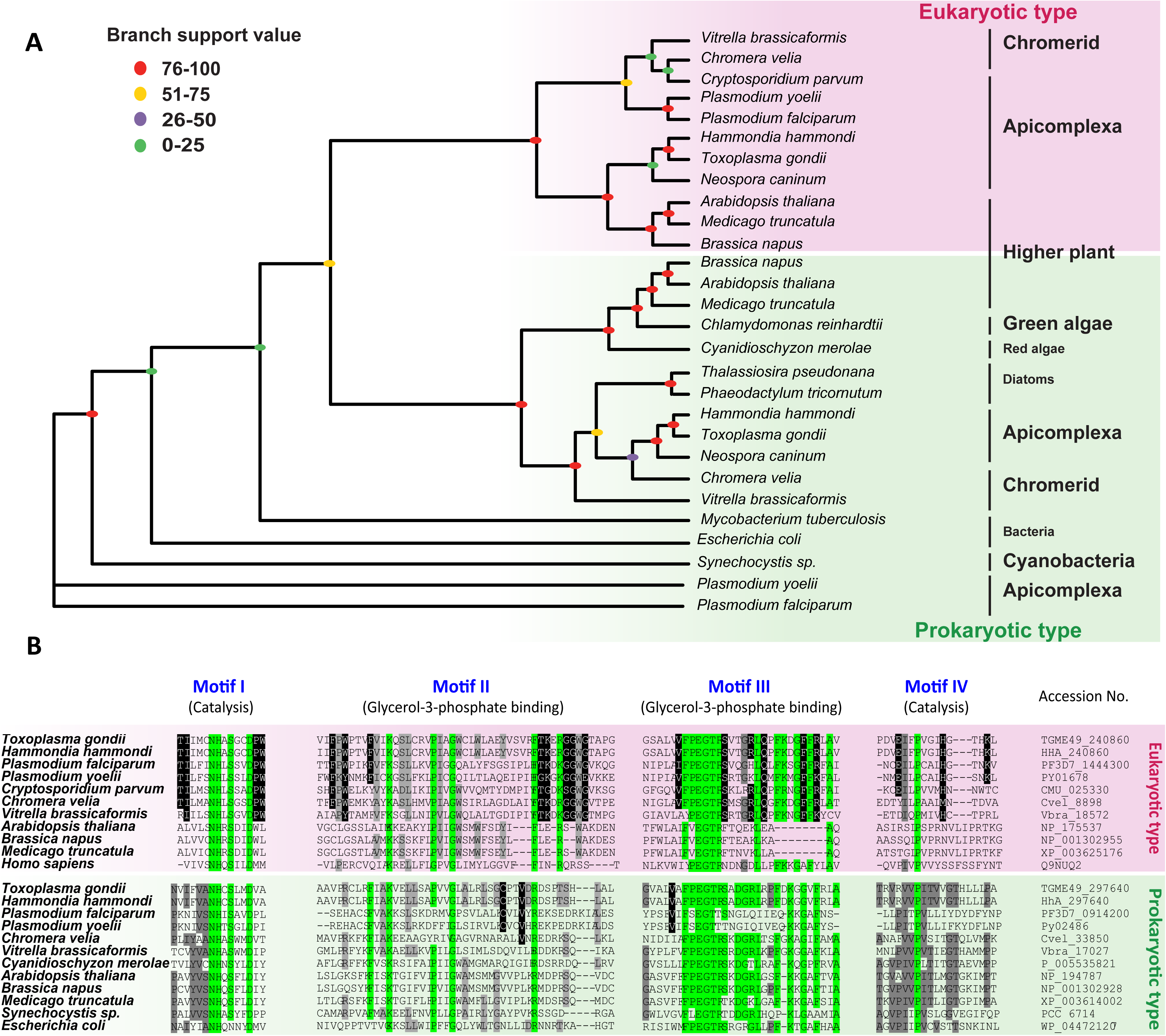
**(A)** Phylogenetic analysis of *Tg*AST2 and *Tg*AGPAT and orthologs from various organisms. Maximum likelihood phylogenies of prokaryotic and eukaryotic acylglycerol acyl transferase activity of 28 enzymes. Branch support values are indicated for each node in different colours (0–25, purple; 25–50, green; 50–75, orange; 75–100, red). **(B)** Multiple protein alignment of the putative orthologues of *Tg*ATS2 from plant chloroplast and *Tg*AGPAT from ER. Green: identical amino acids (aa) residues of active domain (Motif I-IV), grey: aa residues similar among each clade; black aa conserved in Alveolate enzymes in both clades.

**Figure 2:**
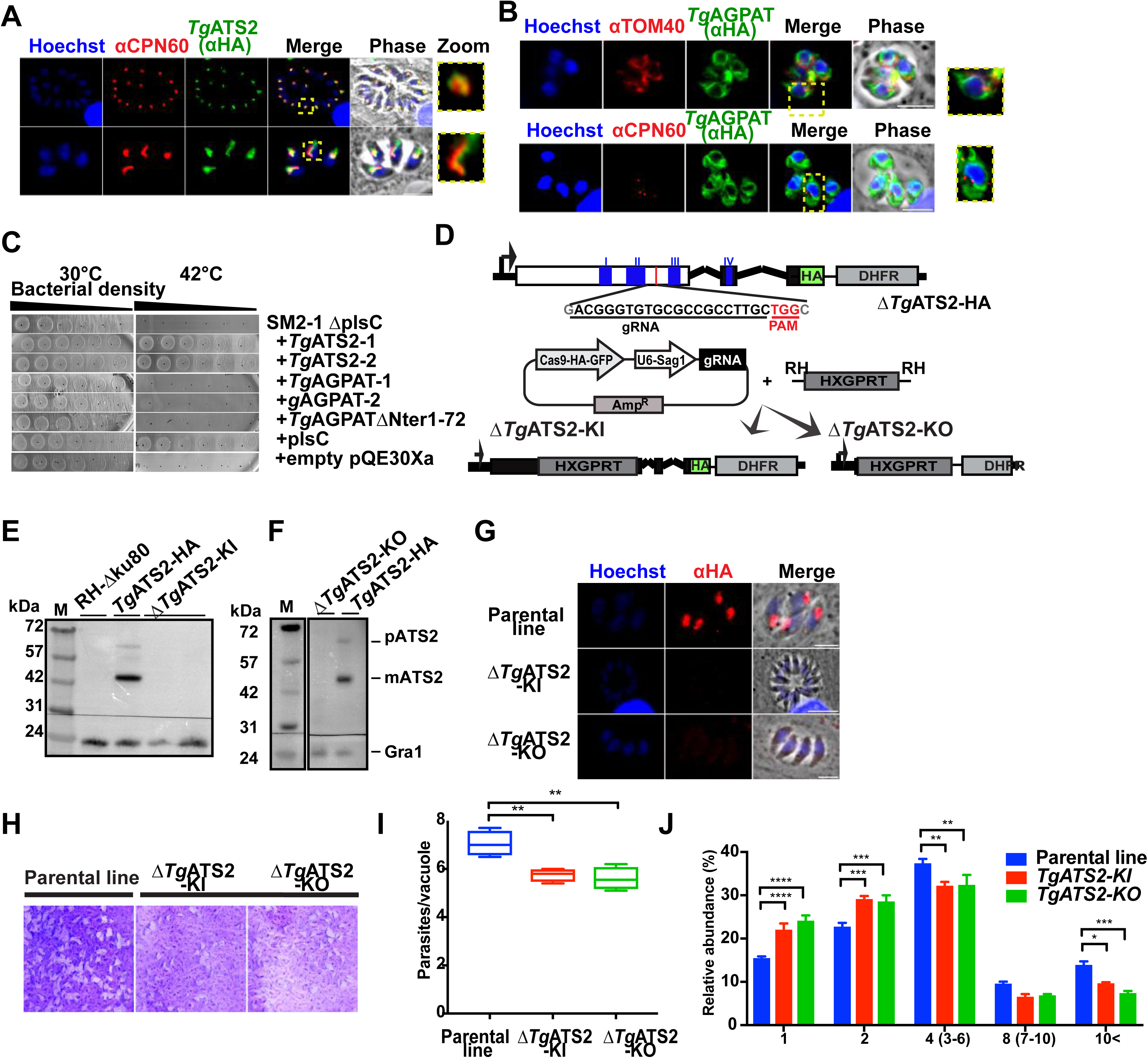
**(A, B)** IFA of stable *Tg*ATS2-HA expressing parasites **(A)** and transient *Tg*AGPAT-HA expression **(B)**. CPN60: apicoplast marker;TOM40, mitochondrial marker. Scale bars=2 μm **(C) Expression of *Tg*ATS2 and *Tg*AGPAT in LPAAT deficient *E. coli* strain SM2-1** SM2-1Δ*plsC* E.coli mutant transformedwith *Tg*ATS2 (1, 2), *Tg*AGPAT (1, 2), TgAGPAT_ΔNter1-72_, *Ec*plsC or empty pQE30Xa expression vector were grown at 30°C (permissive)_ or 42°C (non permissive) for 20 h(n=3). **(D)** CRISPR strategy to generate Δ*Tg*ATS2 mutant lines via integration of HXGPRT cassette within *Tg*ATS2 ORF (knock in; KI) and by *Tg*ATS2 gene replacement by a HXGPRT cassette (knock out; KO) in *Tg*ATS2-HA strain. **(E, F)** Confirmation of *Tg*ATS2-HA loss by Western blot analysis in *Tg*ATS2-KI **(E)** and *Tg*ATS2-KO **(F)** using anti-HA (anti-Gra1: loading control). **(G)** Confirmation of *Tg*ATS2-HA-signal loss in Δ*Tg*ATS2 by IFA using anti-HA (Scale bars=2 μm). **(H)** Plaque assay showing a mild growth defect in Δ*Tg*ATS2 mutants **(I)** Cell-based growth fitness assay confirmed the growth defect in the Δ*Tg*ATS2 mutants 30 h post-infection (n=3). **(J)** Proliferation assay confirmed a replication defect in Δ*Tg*ATS2 mutants (n=3).

To test if *TgATS2* and *TgAGPAT* have predicted 1-acylglycerol-3-phosphate acyltransferase activity, we complemented an *E. coli* temperature sensitive mutant SM2-1 Δ*plsC* lacking the AGPAT activity (Coleman, 1990), with recombinant *TgATS2 and TgAGPAT*. All transformants grew at the permissive temperature of 30°C (**Fig. 2C**). Only those complemented with bacterial *Ec*PlsC and *Tg*ATS2 grew at the non-permissive 42°C (**Fig. 2C**). The constructs with *TgAGPAT*, with or without its long hydrophobic N-Ter extension, did not grow at 42°C likely because of *Tg*AGPAT eukaryotic origin (**Fig. 1AB**). Indeed, eukaryotic AGPATs favour acyl-CoA substrates over acyl-ACP substrates used in bacterial and plastid systems. Thus, it is unclear from our results if *Tg*AGPAT has acyltransferase activity, but *Tg*ATS2 complements PA defective *E.coli* SM2-1 AGPAT enzymatic activity *in vivo*, confirming PA synthetic capability.

To investigate the importance of apicoplast *Tg*ATS2 during tachyzoite life stages, the *TgATS2* locus was disrupted to generate a knock-in (KI) and a knock-out (KO) mutant of *Tg*ATS2 using CRISPR-Cas9 strategies (**Fig. 2D**). Loss of the protein product was confirmed by western blot (**Fig. 2EF**) and IFA (**Fig. 2G**). We attempted to generate a Tet inducible mutant of *Tg*AGPAT but were unsuccessful. We then attempted to knock out *Tg*AGPAT-HA using CRISPR-Cas9 (**Fig. S1**) in either wild type (Δ*Tg*AGPAT) or Δ*Tg*ATS2 (Δ*Tg*AGPAT-Δ*Tg*ATS2) genetic backgrounds, but the parasites were not viable suggesting that *Tg*AGPAT is indispensable, consistent with a genome wide knock-out screen (Sidik et al., 2016). Both Δ*Tg*ATS2 mutants were viable but plaque and replication assays revealed that Δ*Tg*ATS2 had a mild yet significant growth defect with significantly more small (2-4 parasites) vacuoles, and significantly fewer large vacuoles (especially >10parasites that were halved) (**Fig. 2I-K**). Parasite egress was significantly affected in the Δ*Tg*ATS2 mutant (**Fig. S1**) but invasion ability showed no difference with the parental line (**Fig. S1**). Morphology of different intracellular tachyzoite organelles showed no obvious defects except the apicoplast displaying a biogenesis defects (**Fig. S1**).

We further explored parasite replication by assessing Inner Membrane Complex (IMC) formation in Δ*Tg*ATS2 by expressing *Tg*MORN1-mCherry as a marker of the basal complex (Lorestani et al., 2010). The expression of *Tg*MORN1-mCherry in both parental and Δ*Tg*ATS2 lines seemed to show correct localization of *Tg*MORN1 to the ring structure at the basal complex and centrocone (Lorestani et al., 2010). However, careful observations showed an enlargement of residual body (**Fig. 3A**). We further performed IFAs using GAP45 antibody as a marker for the IMC. Indeed, Δ*Tg*ATS2 parasites often displayed a larger area at the centre of big vacuoles (>4 parasites), corresponding to an enlarged residual body (**Fig. 3B**). Statistical analysis confirmed that the size of residual body area significantly increased in Δ*Tg*ATS2 mutant compared to the parental line, especially in large vacuoles (**Fig. 3C**). Interestingly, egressed extracellular parasites often remained tethered at their basal pole via a plasma membrane (PM) structure (**Fig. 3D**). Electron microscopy (EM) revealed that the Δ*Tg*ATS2 parasites, despite normal organelle morphology, incur numerous segregation defects. Most strikingly, Δ*Tg*ATS2 cells frequently showed a late cytokinesis defect in which the dividing cells were tethered to each other at their basal ends (**Fig. 3E**). Parasites were attached through their PM, which was found stretched, forming cytosolic bags likely explaining the enlargement of residual body sizes. The IMC of the affected parasites was fully formed in each daughter cell except at the basal site of daughter cell separation (**Fig. 3E^1,2^**). Furthermore, affected vacuoles displayed enlarged residual bodies that often contained various organelles—including the nucleus, mitochondrion, acidocalcisome vesicles, and other cytosolic material—that appeared to be ejected from the dividing cells due to improper segregation (**Fig. 3E^3,4^**). Pieces of mitochondria were a particular feature within enlarged residual bodies (**Fig. 3E^5,6^**).

**Figure 3:**
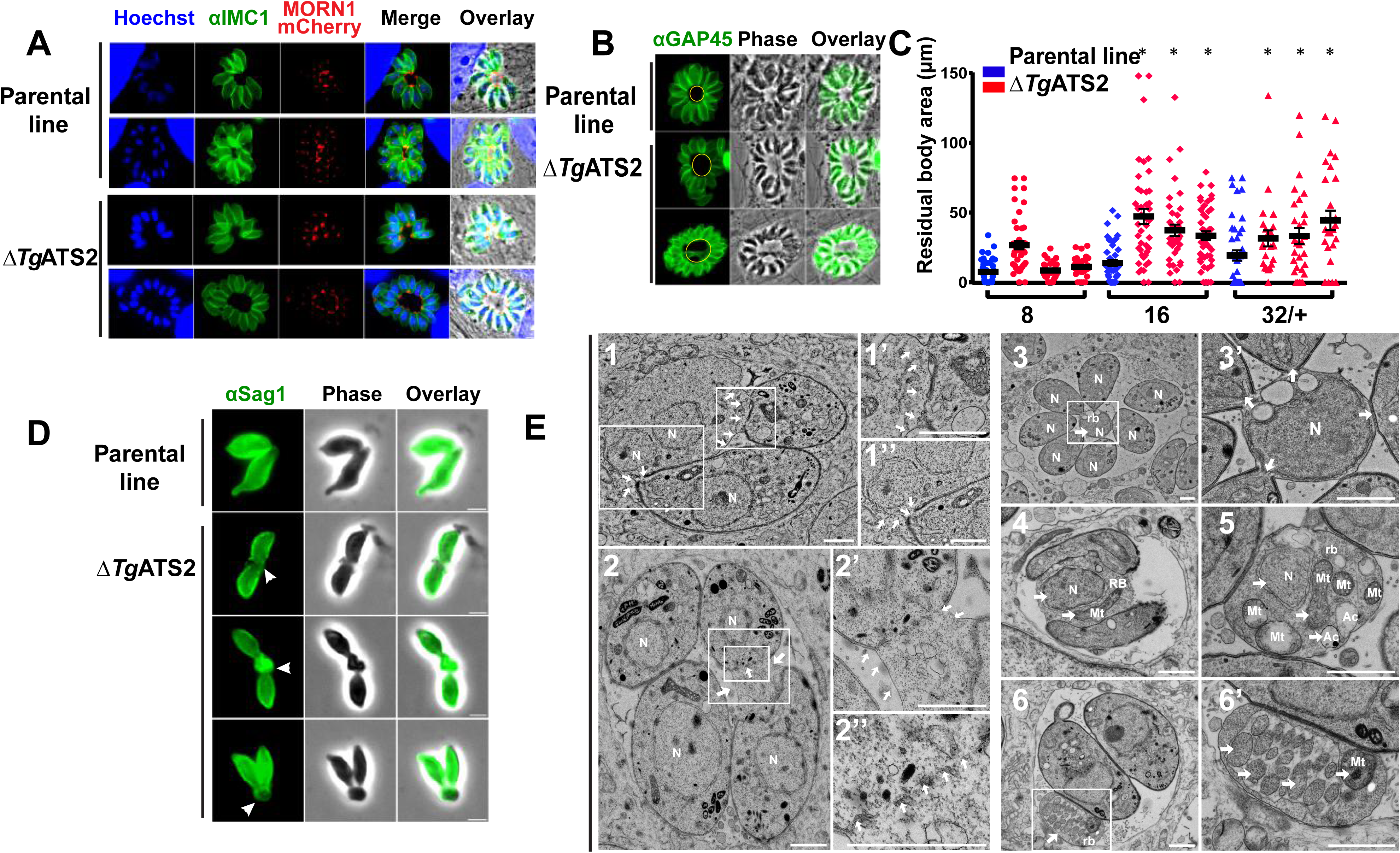
**(A) IFA** of Δ*Tg*ATS2 and parental line transiently expressing MORN1-mCherry (IMC basal tip) and anti-IMC1 (scale bars=2 μm). Confirmation of enlarged residual bodies in Δ*Tg*ATS2 by IFA using anti-GAP45 (IMC marker) **(B)**, and by statistical analysis of residual body size **(C)** (*:*p value* ≤*0.05)* **(D)** IFA observation of extracellular parasite using anti-SAG1 (white arrowhead: PM tether). **(E)** Electron microscopy image of Δ*Tg*ATS2 mutants reveals important cytokinesis defects: major enlargement of the PM, defects in mother cell membrane constriction and cell daughter attachment at the basal pole (**1, 2, enlarged in 1’, 1’’, 2’**, white arrows), IMC fragmentation at the separation sites between dividing parasites (**2’’**). Residual bodies containing unevenly separated nuclei (**1, 3, 3’, 4, 5**), mitochondria (**4, 5, 6, 6’**) and acidocalcisomes (**5**). **N:** nucleus; **Mt:** mitochondria; **rb:** residual body; **Ac:** acidocalcisome (scale bars= 1 μm)

### *Tg*ATS2 disruption dramatically reduces C14:0 FA incorporation into *T. gondii* lipids, skews the LPA/PA ratio, and alters phospholipid abundance and composition

To investigate the role of *Tg*ATS2 in lipid metabolism, we performed lipidomic analysis on the Δ*Tg*ATS2 mutant. Disruption of *Tg*ATS2 resulted in a large and significant reduction of the relative amount of C14:0 (14 carbon FA chain with no desaturation), which is mainly produced via the apicoplast FASII (**Fig. 4AB**). Smaller but significant decreases in C18:1 and C20:1 were also observed (**Fig. 4AC**). In contrast, there was a clear increase in the abundance of C18:0, C18:1(trans), C22:1, C24:1, C20:4, C20:5 and C22:6 (**Fig. 4A-C**), which are most likely scavenged from the host, especially C20:4 and C20:5 as confirmed by analysis of the HFF/FBS content (Welti et al., 2007) (**Fig. S2**). Comparison of the relative FA abundance between Δ*Tg*ATS2 and its parental line showed significant decrease of C14:0, C18:1(cis), C20:0, C20:1 and C22:2 (**Fig. 4C**). These results prove that in addition to the abovementioned cytokinesis defect, the Δ*Tg*ATS2 mutant has a highly modified lipid content that relies less on short-chain, apicoplast-generated FAs and more on long chain fatty acids scavenged from the host (**Fig. 4A-C**).

**Figure 4:**
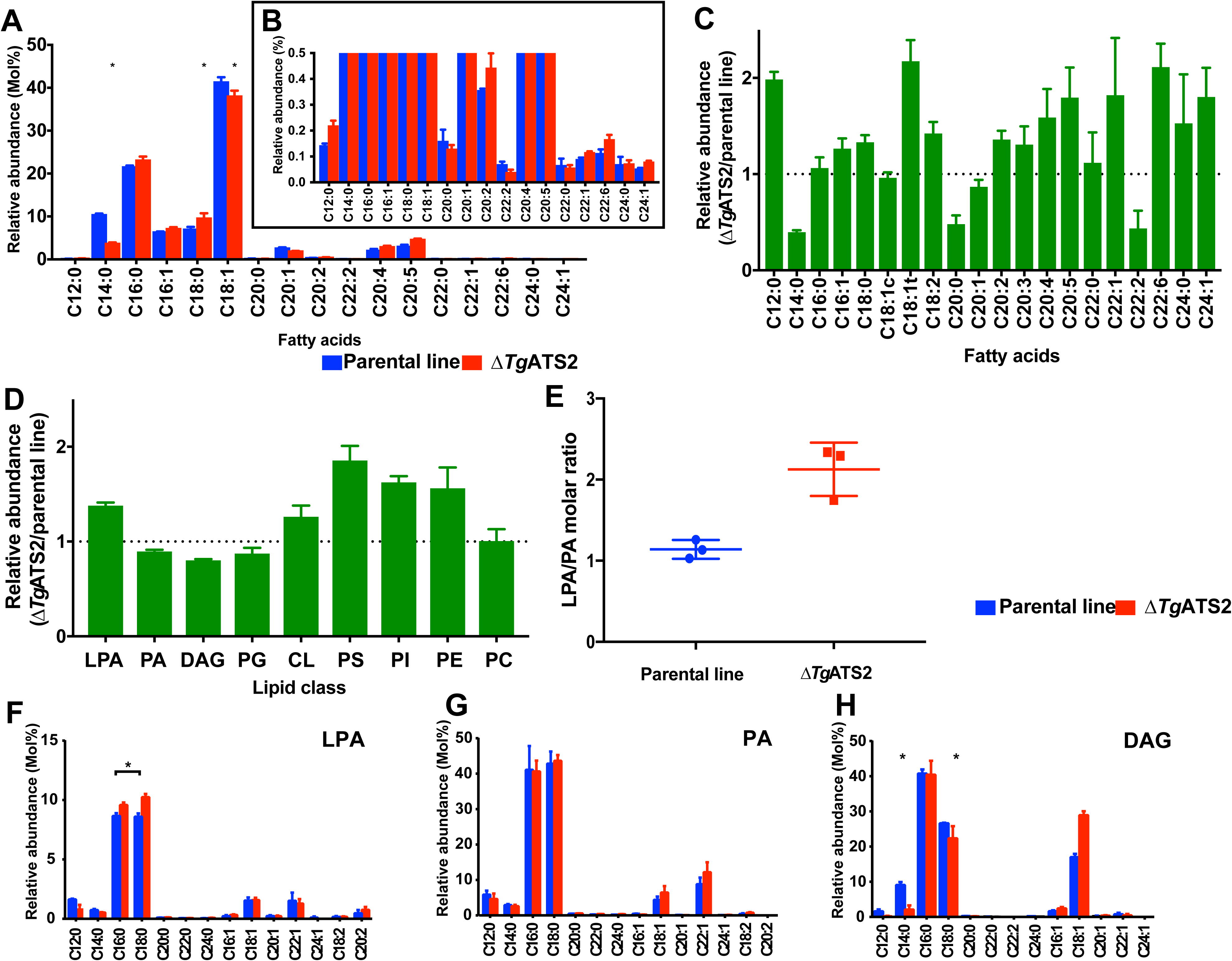
Lipidomic analysis Δ*Tg*ATS2 mutant. **(A, B)** Fatty acid composition of total lipid extracted after 72 h post-infection **(C)** Relative fatty acid abundance of Δ*Tg*ATS2 to the parental line. **(D)** Relative major phospholipid abundance of Δ*Tg*ATS2 to parental line **(E)** LPA/PA ratio. Individual molecular species of LPA **(F)**, PA **(G)**, DAG **(H)**. Fatty acids are shown as Cx:y as x for number of carbons and y for the number of unsaturations. n=4, * indicates *p value* ≤*0.05*

To further investigate Δ*Tg*ATS2 lipid defects, we analysed and quantified each PL class and its FA content. The Δ*Tg*ATS2 mutant accumulates significantly more LPA compared to the control parental line and slightly yet significantly less PA (**Fig. 4DE**), consistent given that LPA and PA are the substrate and product of ATS2 respectively (**Fig. 2C**). This slight difference in PA likely indicates that *Tg*ATS2 is not responsible for the bulk PA synthesis but rather for a specialist function. Importantly, the LPA/PA ratio was drastically affected in Δ*Tg*ATS2 (**Fig. 4E**). We investigated other related PL, namely diacylglycerol (DAG), PC, PE, PI, PS, phosphatidylglycerol (PG) and cardiolipin (CL, **Fig. 4D**). The relative abundance of both DAG and PG significantly decreased in Δ*Tg*ATS2 (**Fig. 4D**). This is relevant since DAG is a direct product of PA, and PG is the sole phospholipid made from PA in plant chloroplasts (Ohlrogge and Browse, 1995). In contrast, the relative abundance of PS, PI and PE increased in the mutant (**Fig. 4D**).

We then examined the FA profiles of each of these lipid classes. LPA had small but significant increases in the amounts of C16:0 and C18:0 in Δ*Tg*ATS2 parasites, whereas small, but significant, decreases in the apicoplast-specific FA, C12:0 and C14:0 were observed in the mutant (**Fig. 4F**). No major difference was observed in PA composition in Δ*Tg*ATS2 parasites (**Fig. 4G**). Strikingly though, DAG, PC, PI and PE all had significantly reduced C14:0 content (**Fig. 4D, Fig. S2**), which is the main product of FASII, and is used by *Tg*ATS1 for bulk *de novo* synthesis of PC, PI and PE (Amiar et al., 2016). This indicates that *Tg*ATS2 likely uses apicoplast generated C14:0 as its major substrate to make these lipids. In contrast, the levels of two long, polyunsaturated FAs (PUFA), C20:4 and C20:5, in all three major PL (PC, PI and PE) were significantly increased in the Δ*Tg*ATS2 line (**Fig. S2**), which is again consistent with mutant parasites compensating for the lack of *de novo* made FAs by increasing scavenging long chain FAs from the host. The FA composition of foetal bovine serum (FBS) and host cell (human foreskin fibroblast, HFF; **Fig. S2**) confirmed that C18:0 and PUFA (such as C20:4, C20:5) were all found in FBS and HFF cells. This is consistent with the Δ*Tg*ATS2 mutants increasing scavenging of these PUFAs to make PC, PI and PE and likely not fully synthesising them *de novo*. We complemented Δ*Tg*ATS2 and wild type parasites (**Fig. S2**), using exogenous PA(14:0;14:0) the putative product of *Tg*ATS2 and PA(16:0;18:1) made of host derived FAs. Proliferation assays showed that both exogenous PA sources could significantly boost parasite growth (**Fig. S2**), but could not rescue Δ*Tg*ATS2 growth phenotype. This indicates that the PA source needs to be made *de novo* via *Tg*ATS2 for proper division. Since parasites are capable of scavenging lipids from the host and medium, we determined whether the Δ*Tg*ATS2 imported more PA, and used PC as a control. Δ*Tg*ATS2 imported significantly more PA and PC than the parental control line (**Fig. S2**). Together these data on extracellular Δ*Tg*ATS2 corroborate our lipidomic analyses (**Fig. 4A**) indicating that the mutant scavenges more lipids to compensate for reduced *de novo* synthesis.

### Disruption of *Tg*ATS2 induces a mislocalisation of the parasite DrpC which perturbs parasite cytokinesis, IMC formation and plasma membrane stability

Lipidomic analyses revealed a drastic LPA/PA imbalance in the Δ*Tg*ATS2 mutant (**Fig. 4E**). LPA and PA have important structural influences on membrane architecture by inducing strong local membrane curvatures, which can impact the recruitment and functions of specific dynamins at precise membrane domains for organelle/vesicle fission: synaptic vesicles transport between neurons requires a protein complex composed of a dynamin and an endophilin that exerts an ATS2 activity to create the proper membrane groove where the dynamin can pinch and release the synaptic vesicle; or even in human mitochondrial fission by the protein Dynamin-like 1, HsDrp1, which require insertion, recruitment and regulation through PA (Adachi et al., 2016; Schmidt et al., 1999). In *T. gondii*, there are three known Drps: *Tg*DrpA, *Tg*DrpB, and *Tg*DrpC. TgDrpA and TgDrpB have roles in apicoplast fission, and secretory organelle biogenesis, respectively (Breinich et al., 2009; van Dooren et al., 2009). *Tg*DrpC was recently localized to the basal poles of dividing daughter cells (Heredero-Bermejo et al., 2017). We generated a parasite line expressing *Tg*DrpC fused to a 3xHA tag under the control of its endogenous promoter using CrispR-Cas9 (**Fig. S3**) and localized *Tg*DrpC-HA during the tachyzoite intracellular division cycle in Δ*Tg*ATS2 and its parental line (**Fig. 5A**). In parental line parasites *Tg*DrpC-HA clustered in small punctate-like compartments in the apical post-Golgi area during interphase (**Fig. 5A**). During daughter budding, *Tg*DrpC re-localized to form two distinct ring-like structures coinciding with the growing ends of the IMC from the budding daughter cells which constricted at the base of the mother cell during cytokinesis and eventually formed basal caps on the each newly divided parasite (**Fig. 5AB**).

**Figure 5:**
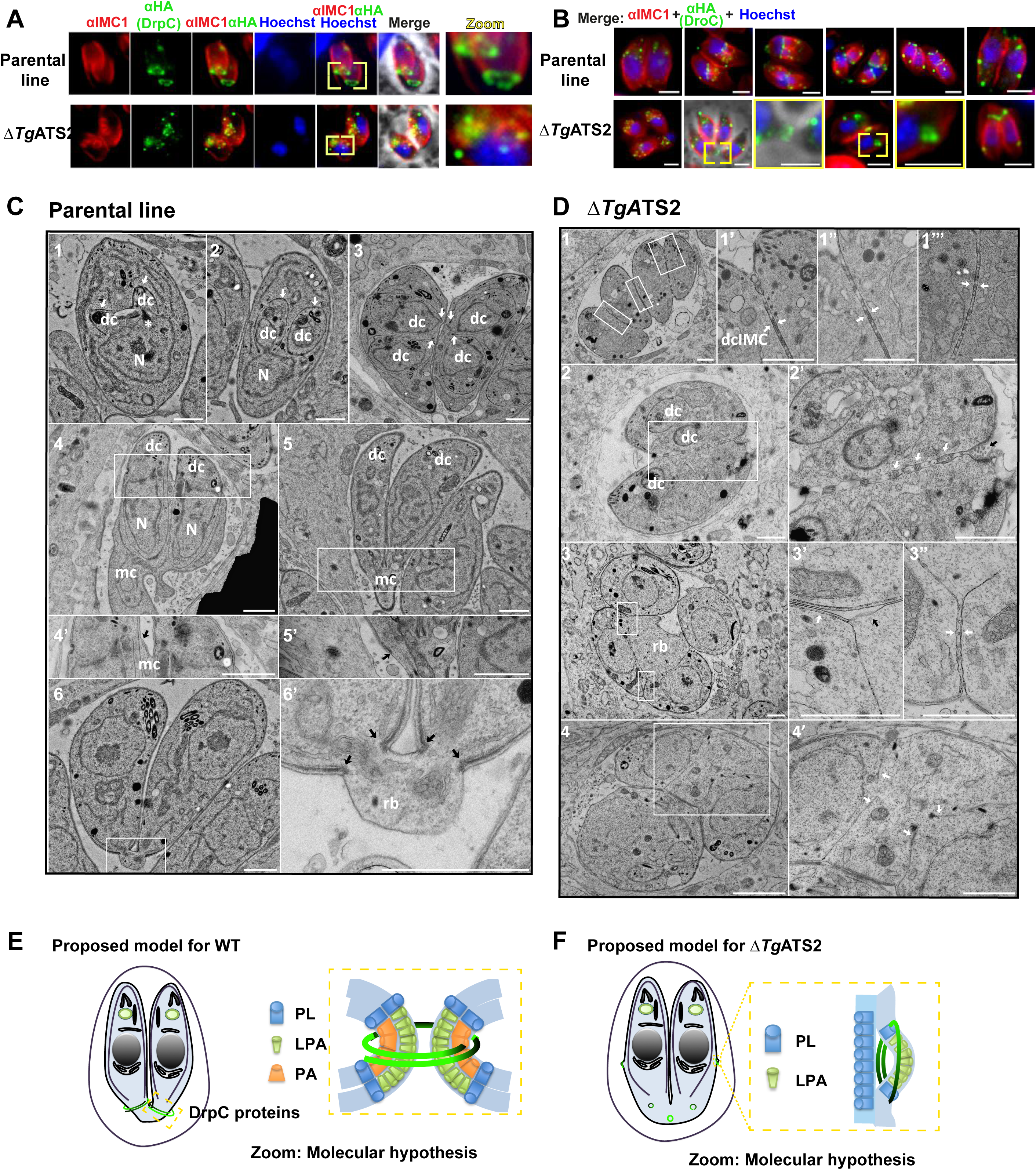
Δ*Tg*ATS2 induces the mislocalization of *Tg*DrpC and cytokinesis defects during tachyzoite division. **(A)** IFA localization of *Tg*DrpC-HA expressed in parental line shows ring structures at the growing ends of daughter cells during division (top panel) but fails to do so when expressed in Δ*Tg*ATS2 (bottom panel). **(B)** *Tg*DrpC-HA localisation during tachyzoite division cycle in the parental line (top panel) and its mis-localization in Δ*Tg*ATS2 mutant. Scale bars= 2 μm **(C, D)** Electron microscopy observation of basal pole of IMC parental line **(C)** and ΔTgATS2 **(D)**. **(C^1-3^)** Endodyogeny starts with the formation of daughter cells (dc) by growth of IMC (white arrows) and organelles segregation. IMC scaffolding then grows towards the basal pole (white arrows) encompassing divided organelles (eg nucleus N). **(C^4-5’^)** Recycling and biogenesis of PM (black arrow) ends daughter cell emergence from mother cell (mc). **(C^6-6’^)** Division ends by cytokinesis through constriction of both IMC and PM at basal pole (black arrows) to form a small residual body (rb). **(D)** Δ*Tg*ATS2 shows an incomplete separation of daughter cells during cytokinesis with absence of PM biogenesis between closely apposed IMC **(1,3 and insets, white arrows)**, presence of vesicle/cisternae inside membrane structures at the inter-IMC space **(2’ white arrows)**, absence of mother IMC **(3’ black arrow)**, absence of basal constriction forming large residual bodies leaving floating daughter IMC **(3,4, 4’ white. arows)**. Scale bar = 1μm. **(E,F)** Proposed molecular model for *Tg*DrpC function during endodyogeny and cytokinesis in WT parasite (E) and Δ*Tg*ATS2 (F) LPA and PA molecules induce positive and negative curvature creating grooves in biological membranes for *Tg*DrpC to insert at specific sites during division and exert its pinching function during endodyogeny **(E)**.

In Δ*Tg*ATS2, localization of *Tg*DrpC-HA was drastically affected only during division (**Fig. 5B**). Whilst interphase punctate localisation was unchanged, *Tg*DrpC-HA failed to form the typical ring structures at the initiation and during division. Instead, it scattered in the cytosol, or formed rings pushing on the side of mother IMC, or not constricted at the daughter basal pole. To investigate the reason of the specific mislocalisation of TgDrpC after TgATS2 disruption, we performed *in silico* analyses, which showed that (i) TgDrpC is the closest homolog to the human Dynamin-like 1, HsDrp1, which allows mitochondrial fission through its interaction with PA via its Stalk domain including a loop with specific hydrophobic residues (Adachi et al., 2016; Adachi et al. 2017), (ii) that both the Stalk domain and this PA binding loop are conserved in TgDrC, (iii) but are absent in the other T. gondii TgDrpA, and TgDrpB (**Fig. S3**). To confirm this, we tagged and monitored the localization of other TgDrps in the Δ*Tg*ATS2 background. No obvious change in localization of *Tg*Drp*A* was observed in Δ*Tg*ATS2 parasites, even during the fission of the apicoplast (**Fig. S4**). Similarly, no change in the localization of *Tg*DrpB was observed upon *Tg*ATS2 disruption as (not shown), as expected from both IFAs and EM, showing no defect in rhoptries biogenesis, where TgDrpB is involved. confirming protein sequence and domain analysis. Taken together, our data strongly suggest that perturbation of the LPA/PA content in Δ*Tg*ATS2 corrupts the orderly processes of cell division by inducing mis-localization of *Tg*DrpC.

In dividing parasites, it is commonly seen that the PM is kept connected between two recently divided cells so that the cells are stuck together and distributed to daughter cells during cytokinesis. Very little is known on the molecular mechanisms of the PM segregation/biogenesis during cytokinesis. In the parental line parasites, initial steps of endodyogeny showed the formation of the daughter cell apical pole along with organelle division before the formation of the daughter cells within the mother cell (**Fig. 5C**). Emergence of the daughter cells initiates the apical-to-basal biogenesis of their PM, partly recycled from the mother (**Fig. 5C^1^-^5^**), and ends by a constriction of both IMC and PM at the basal poles, leaving a small basal residual body (**Fig. 5C^6^**). In contrast, there were many division and cytokinesis defects in Δ*Tg*ATS2, which were unable to separate although new round of daughter formation could be initiated (**Fig. 5D^1^**). Daughter cells were found tightly apposed at normal emergence sites and their PMs were often missing between daughter IMCs. Instead, interconnection of PM, vesicles or cisternae could be observed at these apposition sites and at the basal end of dividing cells (**Fig. 5D^2^**). These defects suggested issues at the PM composition and/or problems in membrane fusion/fission sites. Furthermore, there was no constriction of both IMC and PM from daughter cells resulting in enlarged residual bodies containing organelles and cytosol portions (**Fig. 5D^34^**). Altogether, these changes in Δ*Tg*ATS2 likely results from the imbalance of the LPA/PA ratio impacting membrane curvature required during cytokinesis (**Fig. 5EF**)

### Nutrient starvation enhances the synthesis of FA by apicoplast FASII in *T. gondii* and blocks intracellular proliferation of *P. falciparum* blood stages lacking a functional FASII

Since *Tg*ATS2 has a role in maintaining parasite lipid homeostasis, we set out to determine the balance of *de novo* synthesized versus scavenged lipids in Δ*Tg*ATS2 using a stable isotope precursor of apicoplast synthesised fatty acids, U-^13^C-glucose. (Amiar et al., 2016; Dubois et al., 2018; Ramakrishnan et al., 2012). Incorporation of ^13^C within each FA species is detected by increase of mass and determined in relation to non-labelled FA. Distribution of ^13^C incorporation to each FA isotopologue is shown as its own mass (M) plus number of ^13^C carbon incorporation, *i.e.* M+x. In both parental and Δ*Tg*ATS2 mutant lines, we observed significant differences of ^13^C incorporation in C14:0, C16:1, C18:0 and C18:1 (**Fig. 6A**). Isotopologue distribution of apicoplast-signature C14:0 showed that Δ*Tg*ATS2 had ^13^C incorporation up to M+14 but major incorporation occurred at lower masses (M+8, M+10) than the parental (M+12, M+14, **Fig. 6B**). This indicates that the FASII is active in Δ*Tg*ATS2 yet slowed down in the process of making C14:0, thus explaining the C14:0 reduction previously detected (**Fig. 4A**). Similar significant, yet milder, results were observed for C16:0 isotopologue distribution although overall incorporation was similar between parental and Δ*Tg*ATS2 (**Fig. 6C**). C18:0 in Δ*Tg*ATS2 had higher ^13^C incorporation than the parental and its isotopologue distribution showed more short FA from the apicoplast (**Fig. 6D**).

**Figure 6:**
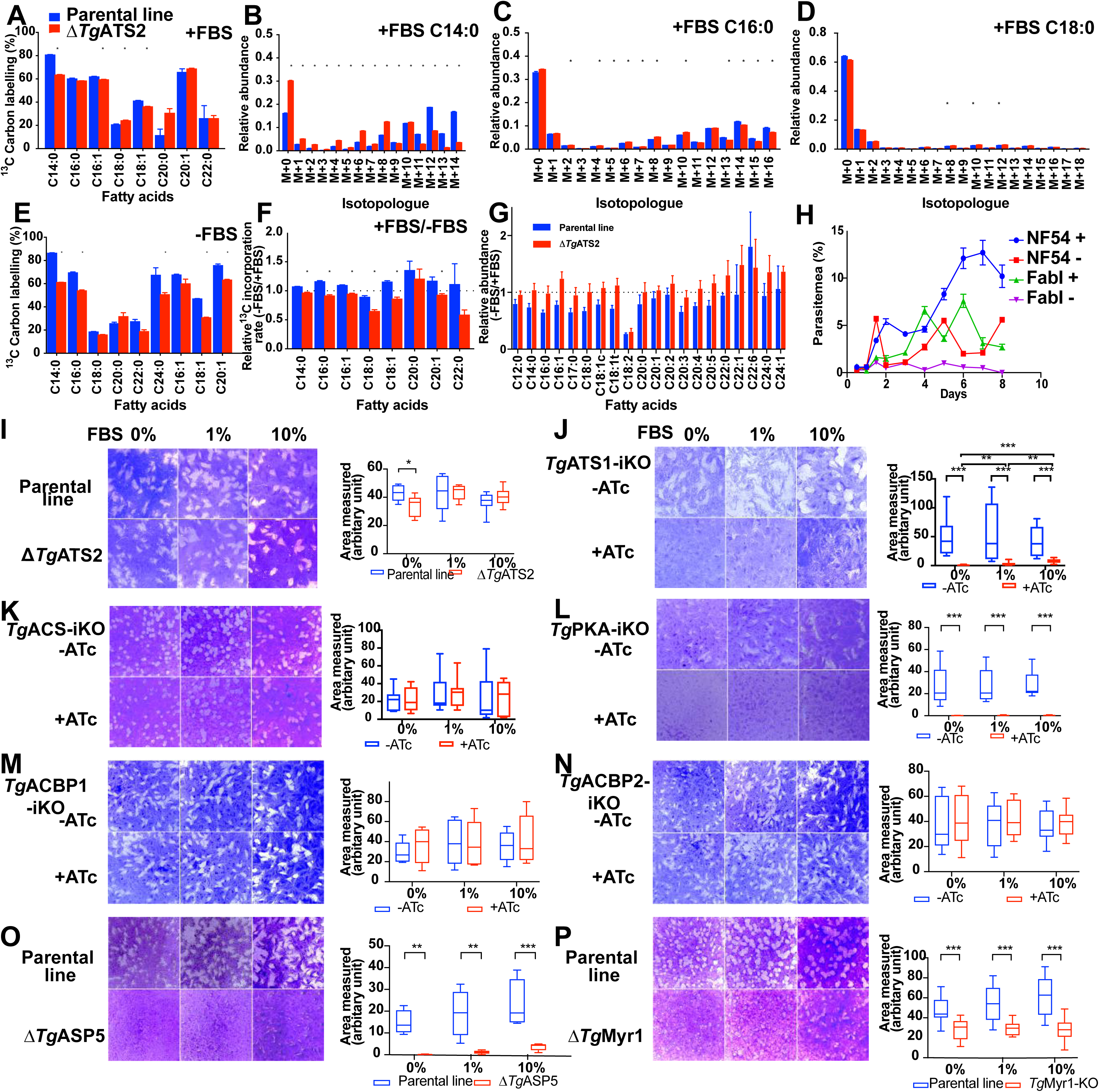
FASII can adapt its metabolic capacities upon changes in nutritional environment in *T. gondii* and is critical during lipid starvation in *P. falciparum* blood stages. **(A-G)** U-^13^C-glucose labelling for 72 h to monitor apicoplast FA synthesis by ^13^C incorporation to each fatty acids (Blue, parental line, Red, ΔTgATS2) **(A)** ^13^C incorporation to each fatty acids in 1% FBS **(B-D)** Mass isotopologue distribution in 1% FBS for C14:0 **(B)**, C16:0 **(C)** and C18:0 **(D)** The x-axis shown as ‘M+X’ represents mass with ‘X’ ^13^C atoms incorporated during the FA synthesis. **(E)** ^13^C incorporation to each fatty acids in 0.2% FBS. FASII metabolic activity increased upon FBS starvation in the parental line but not in Δ*Tg*ATS2.**(F)** Change in the ^13^C incorporation between 0.2%FBS and 1% FBS (-FBS/+FBS) **(G)** The relative abundance of each FA (-FBS/+FBS) **(H)** Asexual blood stage growth assay of *P. falciparum* FabI-KO and its parental line (NF54) in regular (lipid rich) culture medium and lipid starved medium. **(I)** Plaque assays conducted in 0, 1, or 10% FBS in different *T. gondii* mutants and their reference strain; *Tg*ATS2, *Tg*ATS1, *Tg*ACS, *Tg*PKA, *Tg*ACBP1, *Tg*ACBP2, *Tg*ASP5, *Tg*Myr1. n=3, or more. *, p ≤0.05; **, p≤0.01; ***, p≤0.001; ****, p≤0.0001.

Lipidomic analyses thus indicate that both scavenged and *de novo* lipid fluxes are modified in Δ*Tg*ATS2. To tease out the impact of host nutritional environment on both pathways, we sought to measure parasite lipid fluxes under adverse host nutritional/lipid conditions, through limitations in FBS concentrations in parasite culture media. Interestingly, GC-MS analysis revealed that ^13^C incorporation into all FASII-generated and further ER-elongated FA products, i.e. C14:0, C16:0, C16:1, C18:0, C18:1, C20:1, was significantly higher by 5-15% under FBS starvation in the parental line (**Fig. 6EF, S5**). In addition, ^13^C incorporation into most FAs is increased in the wild type parental line (**Fig. 6F**). These results suggest that apicoplast *de novo* FA/lipid synthesis can be upregulated during FBS starvation to compensate for the lack of nutrients in the external environment. However, in Δ*Tg*ATS2, the ^13^C incorporation into each FA was decreased by FBS starvation (**Fig. 6F**). No morphological changes could be observed by IFA in the FBS starved Δ*Tg*ATS2 mutant (Fig.5). Both parental line and Δ*Tg*ATS2 mutant showed a significant reduction in the synthesis of C18:0 in FBS starved conditions, suggesting that C18:0 is predominantly obtained by scavenging from the host cell (**Fig. 6F**). Since availability of lipids from the environment is limited, the fatty acid abundance in the parental line was decreased (**Fig. 6G**). Interestingly however, the fatty acid abundance in Δ*Tg*ATS2 was increased in most of its FA species during FBS starvation (**Fig. 6G**).

Although we observed a defect in the activation of FASII in Δ*Tg*ATS2, Δ*Tg*ATS2 was nevertheless viable during FBS starvation. This suggests that if FASII is active, regardless of the level of FASII activity, the parasites are viable under FBS starvation, consistent with its essential role in tachyzoites. However, in *P. falciparum*, FASII is not essential during nutrient replete blood stage but is activated under lipid starvation, apparently to compensate for reduced availability of scavenge-able lipids (Botte et al., 2013; Yu et al., 2008). Our results in *T. gondii* led us to re-think the current hypothesis regarding the dispensability of the apicoplast FASII in *P. falciparum* blood stages and to test the essentiality of malaria parasite apicoplast FASII under nutrient/lipid starved conditions. We grew *P. falciparum* FASII KO, Δ*Pf*FabI (Yu et al., 2008) and its parental line (NF54) in either regular (i.e. lipid rich) culture medium or in “lipid-starved” medium (Botte et al., 2013; Mi-Ichi et al., 2007). Both NF54 and Δ*Pf*FabI grew normally in the regular culture medium (**Fig. 6H**). In the lipid-starved medium, NF54 was viable but grew significantly slower than in lipid replete conditions whereas Δ*Pf*FabI grew only for the first 2 days in lipid starved media, but after 4 days, a sharp decrease in growth occurred, and this led to a complete loss of detectable parasites after 8 days (**Fig. 6H**). This proves that FASII is required for the malaria parasite to adapt its lipid metabolism in response to an adverse host lipid environment, a similar situation to that revealed here for *T. gondii*.

Since environmental FBS starvation induces an increase of *de novo* lipid synthesis, we investigated the effect the lipid-nutrient depleted conditions (i.e. 0, 1, 10% of FBS) on various mutants involved defective in lipid metabolism in *T. gondii*. We assessed parasite growth by plaque assay and quantified plaque area. The wild-type and parental parasite lines could grow equally well in DMEM supplemented with 0, 1, or 10% FBS (**Fig. 6I-P**), without affecting the integrity of HFF host cells. FBS starvation only affected growth of Δ*Tg*ATS2 mutant under 0% FBS (**Fig. 6I**). FBS availability had a much stronger effect on iΔ*Tg*ATS1-KO (Amiar et al., 2016) **Fig. 6J**). Although *Tg*ATS1-KO already grows much less in the regular culture conditions (*i.e.* 1% FBS), starvation under 0%FBS led to the quasi-absence of plaques whereas an increase up to 10% FBS led to significantly larger plaques than in the 0% and the 1% FBS (**Fig. 6J**). This suggested that in the absence of the major *de novo* phospholipid precursor synthesis pathway, the parasite could compensate the growth defect by accessing more host lipid resources. The acetyl-CoA synthetase, *Tg*ACS (Dubois et al., 2018), was slightly responsive to FBS starvation (**Fig. 6K**). Interestingly, proteins not involved in bulk membrane/lipid synthesis, such as *Tg*PKA-iKO, do not display an aggravated phenotype under nutrient starvation (**Fig. 6L**) (Uboldi et al., 2015).

Since host fatty acid binding proteins (FABP) are upregulated upon tachyzoite invasion (Hu et al., 2017), we searched the genome of *T. gondii* for homologs of FABP that could be responsible for the transport of FAs in the parasite during starvation but found none. Instead, we found two proteins belonging to the closely related family of acyl-CoA binding protein (ACBP), i.e. *Tg*ACBP1 and *Tg*ACBP2. We identified that *Tg*ACBP1 and *Tg*ACBP2 localised at the parasite cytosol and mitochondrion, respectively (**Fig. S6**). Then we generated inducible knock-down parasite lines for both (**Fig. S6**). However, plaque assays showed that both proteins were dispensable during tachyzoite life stages and neither were responding to FBS starvation (**Fig. 6MN**), suggesting that neither *Tg*ACBPs are involved as effectors for the adaptation to nutritional environment.

We the hypothesized that parasite effectors putatively exported into the PVM or towards the host cell could be used by the parasite to collect putative host membrane material generated during FBS starvation. To test this, we investigated *Tg*ASP5, a Golgi resident aspartyl protease that controls the non-canonical trafficking pathway of parasite effectors towards the parasitophorous vacuole and the host cell, during FBS starvation (Bougdour et al., 2014). Strikingly, FBS starvation significantly exacerbated the growth defect in *Tg*APS5 mutants, whereas there was no change the knockdown of TgMYR1, the canonical system to export effectors towards the host (Franco et al., 2016), the under these conditions (**Fig. 6O, P**). Together, these data provide evidence that parasite effectors are trafficked to the host cell solely thought the TgASP5 export pathway to modulate to nutrient starvation to enhance the ability to scavenge needed resources.

### Nutrient depletion/starvation induces the formation of multi-membrane bound vesicles in host cells that are taken up by the parasite

FBS starvation induces parasites to activate an adaptation pathway that apparently includes shifts the balance between *de novo* synthesis and scavenging from the host, as well as scavenging from the host/parasite environment. To investigate potential changes to the host cell and hence host/parasite interactions during lipid starvation we performed EM on starved (0, 1 or 10% FBS) HFF host cells infected with either the parental parasite line or Δ*Tg*ATS2. Growth in 10% FBS led to no obvious phenotype changes in the hosts cells or the parental parasite line or the Δ*Tg*ATS2 mutant (**Fig. 7AB**), but reduction to 1% and 0% FBS induced striking changes in the host cells, which became massively vesiculated irrespective of whether they were infected with the parental line or Δ*Tg*ATS2 (**Fig. 7AB**). Such vesiculation was not observed in uninfected HFF host cells put under nutrient starvation. Giant multi-vesicular bodies (gMVB), i.e. large membrane bound compartments containing various smaller vesicles, were frequent in 1% FBS grown cells (**Fig. 7A^1^-^4^**) and very numerous at 0% FBS (**Fig. 7B^1^-^4^**). The gMVBs could arise from the host ER, as the ER could be seen swelling and forming networks containing large lipid bodies (**Fig. 7B^3^**). gMVBs were also often seen in close apposition/contact with the mitochondria and/or ER network, indicating that material could also be transferred from mitochondria (**Fig. 7B^3,5^**). However, gMVBs were more often observed arising directly from the host nuclear envelope, likely representing a major contributor for their formation (**Fig. 7B^6^-^8^**). The gMVB accumulated in close vicinity with the PVM, which houses the parasite during its intracellular development and serves as the exchange interphase between the host and the parasite. The gMVBs were not only close to the PVM but appeared to be interacting with the PVM with host material and vesicles from the gMVB apparently “percolating” through the PVM (**Fig. 7B^4^**) or directly from their originating organelles (**Fig. 7B^5^**) to eventually be found in the PVM (**Fig. 7A^2^B^1^**). Indeed, these vesicles appeared in both wild-type and Δ*Tg*ATS2, suggesting that the host cell is responding to the nutrient deficiency in a similar manner irrespective of whether it is infected with the parental parasites or Δ*Tg*ATS2. The Δ*Tg*ATS2 parasite cytokinesis phenotype (e.g. **Fig. 3**) was still observed and apparently exacerbated in 0% and 1% FBS growth medium (**Fig. 7B**). This vesicle/gMVB formation and trafficking to and within the PVM was not apparent in high (10%) FBS medium, suggesting that host gMVBs somehow allow the parasite to increase its lipid scavenging in the absence of nutrient rich serum. This is first report of gMVB containing multiple vesicles, that (i) are dependent and induced by nutrient availability, and (ii) originate directly from diverse host organelles. The gMVBs are clearly different from the lipid droplets of host cell origin used as a lipid source by *T. gondii* (Nolan et al., 2017; Romano et al., 2017).

**Figure 7:**
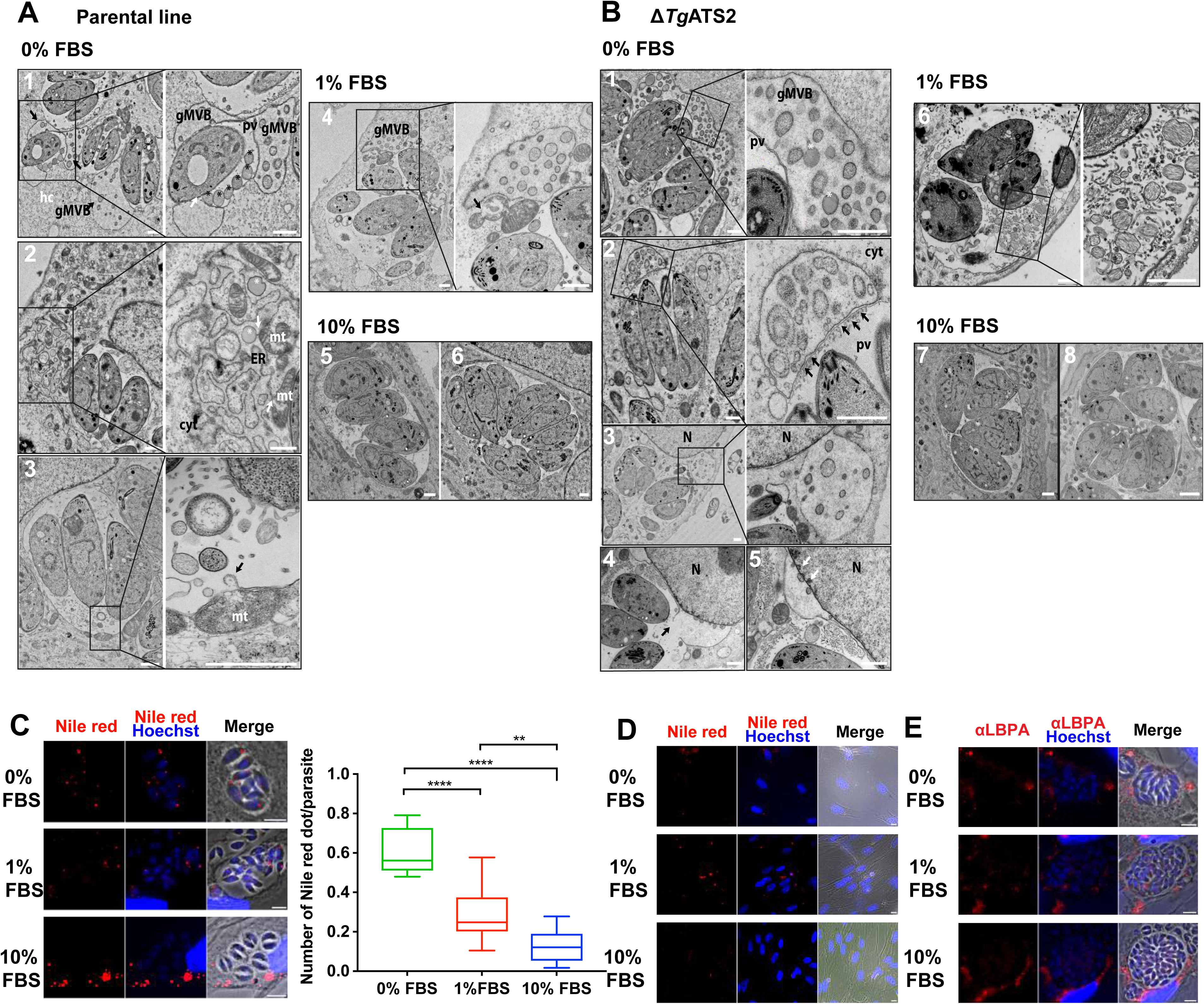
Nutrient starvation unveils formation of multi vesicular bodies from host cell organelles, which content is imported towards parasites. (Transmission electron micrographs of intracellular wild types parasites **(A)** and Δ*Tg*ATS2 mutant parasites **(B)** grown in 0,1 and 10% FBS. Nutrient starvation (i.e. 0 and 1% FBS) induce formation of giant multivesicular bodies (gMVB) in the host cell (hc), containing various vesicles including lipid bodies-like (white stars). In starvation, gMVB localized in the cytosol (cyt) in contact to the parasitophorous vacuole (pv) (**A^1,2,4^B^1,2^**), and their content was imported through and into the PV (**A^1^ black stars, B^2,6^ black arrows**), gMVB were arising from host endoplasmic reticulum (ER, **A^2^**), mitochondria (mt, **A^2,3^**) and mainly swollen nuclear envelope (N, **B^3,4,5^**). 10% FBS did not induce gMVB formation in both parental and Δ*Tg*ATS2. Scale bar= 1μm. **(C)** Nutrient starvation induces a significant increase of lipid droplets within the parasite and its PV as measured by IFA using Nile red (Nile Red dots were counted for 100 or more parasites n=3, **, p≤0.01; ***, p≤0.001; ****, p≤0.0001). **(D)** Nutrient starvation induces a decrease of lipid bodies in uninfected HFF host cells as measured by IFA using Nile red. **(E)** IFA shows that import into parasites of LBPA (anti-LBPA) is not affected by nutrient starvation.

Since lipid droplets from the host can also be used by the parasites as lipid sources, it is possible that FBS starvation leads to increased host cell lipid droplet import. Nile red staining indeed confirmed that FBS starvation induced a significant increase of the amount of lipid droplets into the parasites and its PVM (**Fig. 7C**), In contrast, low FBS content resulted in a reduced amount of lipid droplets in uninfected host cells whilst high FBS content increased their presence in the host cells alone (**Fig. 7D**). This further indicates that increase of import of lipid droplets to the parasite is upregulated by the parasite during FBS starvation.

Since gMVB also seem to arise from host mitochondria, we used an anti-lyso-bi-phosphatidic acid (LBPA) antibody to detect LBPA, i.e. a degradation product of mitochondrial cardiolipin (Kobayashi et al., 1998) that can also be scavenged by the parasite during intracellular development (Romano et al., 2017) (**Fig. 7E**). LBPA was found surrounding the PVM in the host cell, within the PV and the parasite, but its localization and intensity remained unchanged in response to reduced FBS content (**Fig. 7E**). Direct salvage of mitochondrial cardiolipin *per se* might not be the primary up-regulated scavenging pathway during lipid starvation.

## Discussion

We have shown that *Tg*ATS2 is an apicoplast acyltransferase able to esterify FAs on LPA to generate PA, a precursor for a wide range of parasite lipids. Perturbation of *Tg*ATS2 resulted in perturbed lipid fluxes, which impacts LPA/PA lipid balance causing mislocalization of *Tg*DrpC and vesiculation during cytokinesis. Furthermore, parasites exhibit considerable metabolic plasticity on both *de novo* synthesis in the apicoplast, and host modification for membrane scavenging critical for adaptation to nutrient limiting conditions in the host (**Fig. 8**).

**Figure 8:**
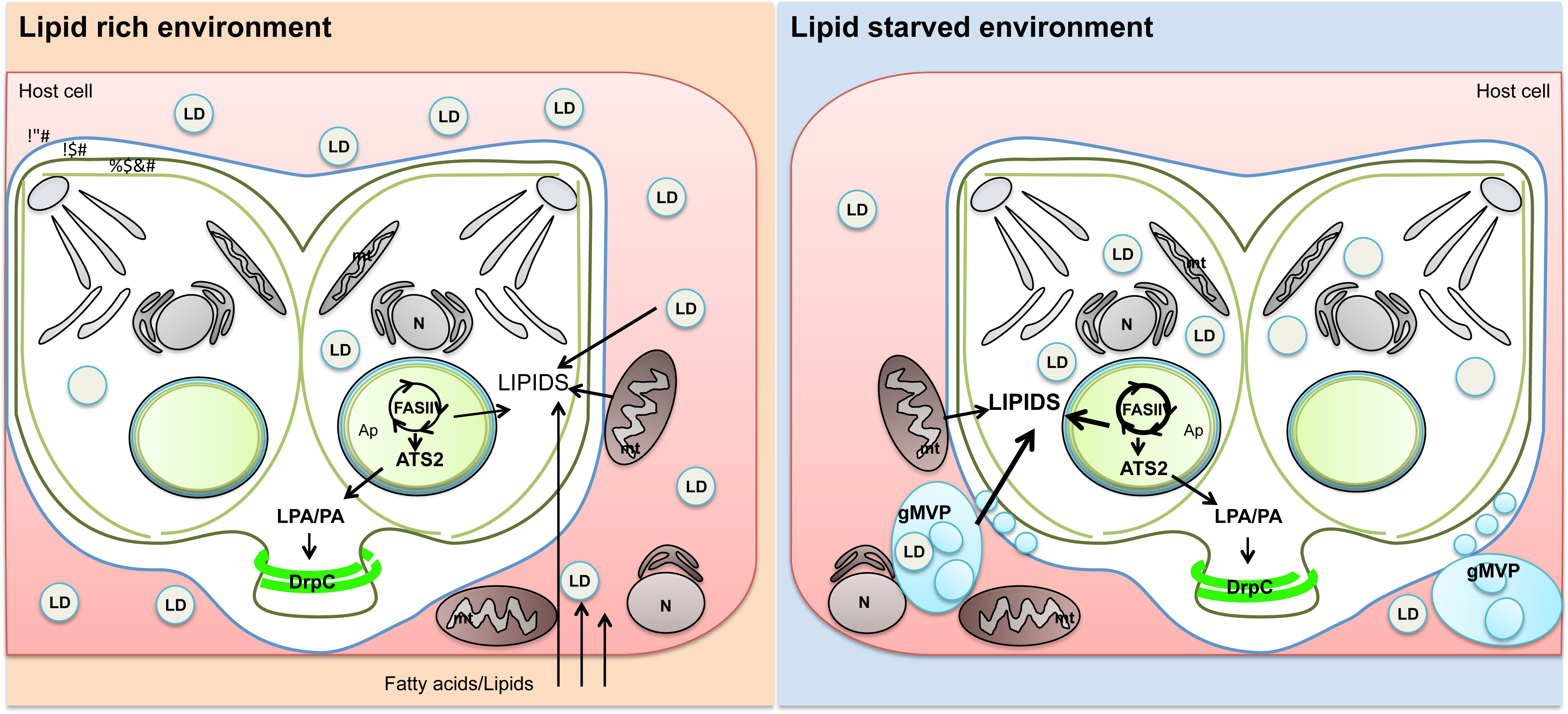
Proposed model for cytokinesis, lipid acquisition and metabolic adaptation under adverse host lipid environment in *T. gondii*. Under lipid rich environment, *T. gondii* can acquire FA and lipids by host cell scavenging, notably from the sequestration of host mitochondria (mt) and lipid droplets but also from the external host cell environment. Under these conditions, the host cell acquires enough nutrients from its external environment to generate numerous lipid droplets and its own lipids. The parasite also needs to generate DA and lipids de novo, notably via the apicoplast (ap) FASII pathway. The apicoplast acyltransferase ATS2 generates PA and regulates the balance of LPA/PA. Such balance creates local membrane curvature to recruit *Tg*DrpC and form plasma membrane, both essential during cytokinesis and division. Under host lipid starved environment, the parasite adapts its metabolism by up-regulating its apicoplast FASII capacities to produce more fatty acid to compensate their absence from the host cell. Concomittantly, the parasite induces morphological changes in the host cell to increase scavenged ressources. Giant multivesicular bodies (gMVB) are being generated from vesiculation of host cell organelles including nucleus (N), ER and mitochondria. These gMVBs are forming in close vicinity to the parasitophorous vacuole membrane (PVM), and their content imported into the PV.

### PA and LPA roles in membrane curvature and cell division

Membrane PLs have different physical shapes according to the relative sizes between the polar head and the fatty acid tails. Most PLs are cylindrical, while PA is cone shaped and LPA adopts an inverted cone shape, thus their insertion into membrane bilayers facilitates curvature and invagination/evagination (Kooijman et al., 2005).

In human cells, PA is implicated in membrane cell curvature during exocytosis and synaptic vesicles formation from neuronal cells (Chasserot-Golaz et al., 2010; McMahon and Gallop, 2005; Schmidt et al., 1999). More particularly, the release of synaptic vesicles between neurons is dependent on the pinching exerted by dynamins that require endophilin1, an ATS2 homolog, as partner to create LPA/PA curvatures (Burger et al., 2000; Shin and Loewen, 2011). Presence of PA in membranes creates electrostatic interaction to improve penetration of a larger part of dynamin into the lipid monolayer (Burger et al., 2000; Shin and Loewen, 2011). Although parasites could proceed with division to a partial extent, *Tg*DrpC requires a specific LPA/PA environment—environment that is altered in Δ*Tg*ATS2. Fairly interestingly, *Tg*DrpC is the homolog of the human Drp1, which also requires the proper LPA/PA interaction to exert its pinching role in human cells (Adachi et al., 2016; Kameoka et al., 2018). Our results thus reveal the previously unrecognised importance of apicoplast maintaining internal lipid homeostasis. Furthermore, the functional role of *Tg*ATS2 for PA synthesis during division provides a mechanism for the long-standing question of why drugs targeting the apicoplast display a secondary cytokinesis defect.

### Environmental and nutritional conditions drive the adaptation of the apicoplast metabolic capacities as well as the scavenging capacities

Importantly, *Toxoplasma* could increase production of FA in the FASII pathway in nutrient-lipid deprived medium similarly to *P. falciparum* (Botté et al., 2011). Hence, apicomplexan parasites show high metabolic flexibility to obtain FA for the major membrane building blocks required for growth as pointed out by recent studies exploring *Plasmodium* survival in nutrient depleted conditions (Brancucci et al., 2017; Mancio-Silva et al., 2017; Zuzarte-Luis et al., 2017). Importantly, our results demonstrate that *P. falciparum* lacking a FASII and grown in lipid-deprived conditions, were unable to properly proliferate, ultimately dying. This suggests that apicoplast FASII is facultative rather than totally dispensable in malaria parasite blood stage and can be activated during lipid starvation to meet needs for phospholipids. This apparent FASII flexibility is consistent with a growing pool of evidence such including the upregulation of FASII in malnourished patients and the essentiality of the central acyl-carrier protein, ACP (Sidik et al., 2016; Zhang et al., 2018), all summarized in **Table 1** which together show FASII is requisite depending on the environment. Therefore, environmental conditions could have important consequences in treating patients. Indeed, if patients are under stress/nutrient deprivation/malnourished conditions, then the FASII pathway could become a secondary target of choice to help eradicate the parasites. This approach is supported by transcriptomics analysis in infected patients displaying an up-regulation of the FASII and the apicoplast acyltransferase PfG3apiGPAt (homolog of *Tg*ATS1, **Table1**, (Daily et al., 2007). Interestingly, FASII remains active during artemisinin induced dormancy in *P. falciparum* blood stages, and maintain parasite growth (Chen et al., 2014), also suggesting that FASII gets actively regulated and could thus be considered a target for combination therapies and during the development of artemisin resistance.. Altogether these data questions whether isopentenyl pyrophosphate (IPP) synthesis is the sole essential function of the apicoplast during *Plasmodium* blood stage (Yeh and DeRisi, 2011). Rather they put the parasite back into its physiological context where nutrient availability and environmental conditions drive the requirement and regulation of a given metabolic pathway. This redefines what we call an essential gene, where phenotypes might only be seen under starvation conditions. An interesting avenue on investigating the parasite metabolic plasticity in response to environmental changes would be to systematically investigate the phenotype of known and uncharacterized proteins putatively exacerbated under lipid deprived conditions.

**Table 1:**
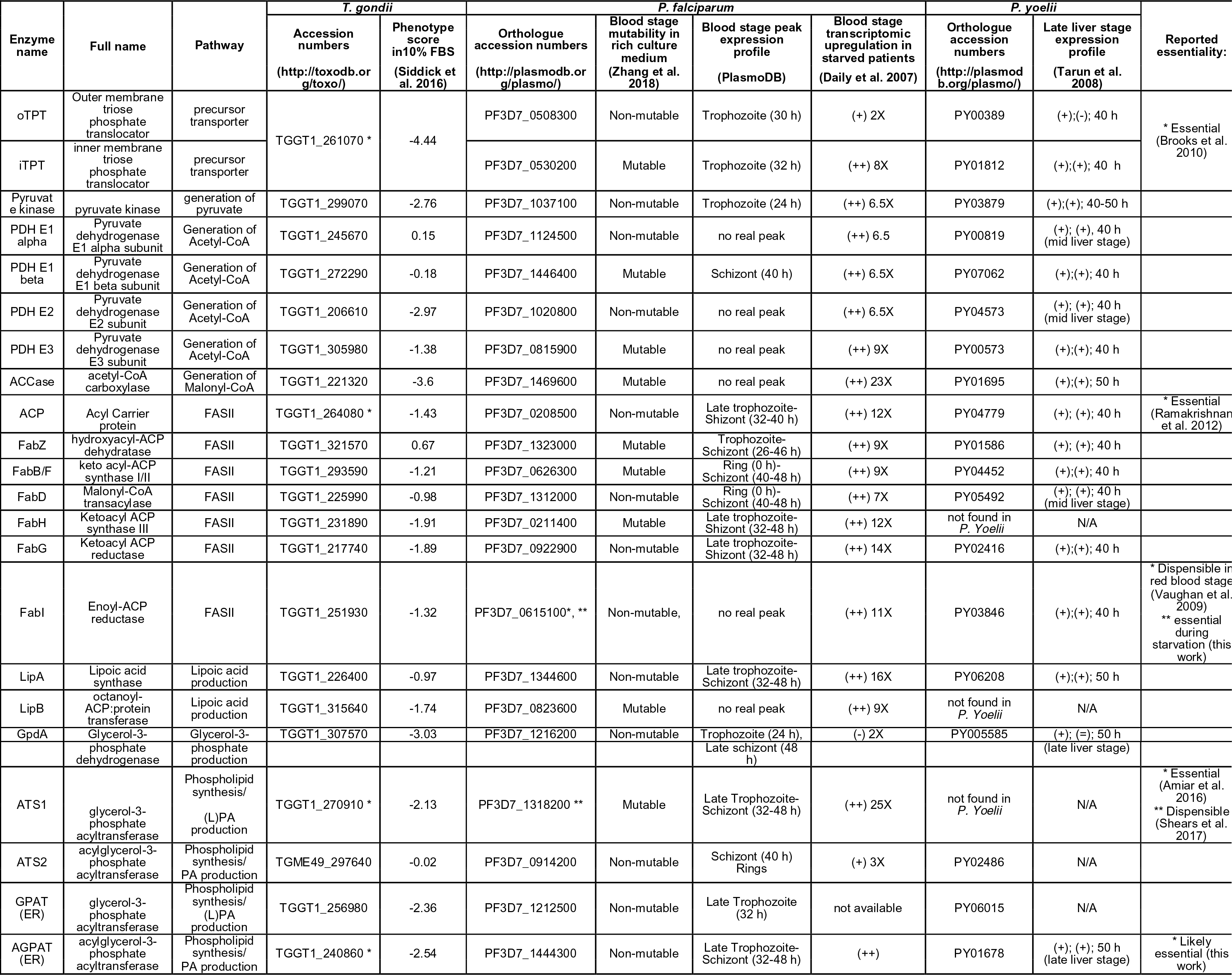
Summary of genomic, transcriptomic and systematic disruption data for genes related to fatty acid synthesis in the apicoplast and phosphatidic acid de novo synthesis. (+) and (++) represent important and very important up-regulation, respectively. (-) stands for downregulation. For transcriptomic analysis analysis: X stands for the approximative fold increase. For Late liver stage expression profile, the table depicts the P. yoelii ortholog number, the regulation of late liver stage (LS) time point 40 h compared to the average blood stage expression level, and the late LS at 50 h compared to average blood stage as per (Tarun et al., 2008). The last time point represents the highest LS expression peak when compared to blood stage.

The increase of scavenging capacities in the Δ*Tg*ATS2 mutant was confirmed via GC-MS analysis of FAs and phospholipid uptake assays, suggesting another sensing-signalling pathway informing the parasite on its own capacities to produce or not the required lipids necessary for intracellular development. While *Tg*ATS2 is probably not the sensor that modulates this, it is clearly in the chain of response important in accelerating growth of the parasite under nutrient deprived conditions, similar to *Plasmodium*. Moreover, EM observation showed unique vesicle structures, gMVB, induced in the host cells upon lipid deprivation that did not correspond to any of the described cellular structures induced by the invasion of *T. gondii.* This suggests that gMVB are actively manipulated and imported by the parasite by the action of parasite effectors sent out to the PV/PVM, via the non-canonical ASP5 pathway. Identification of these effectors warrants future investigations.

A major question raised here is the nature the signalling factor(s) responsible for environmental sensing and metabolic adaption of both apicoplast *de novo* synthesis and scavenging pathways. Both *T. gondii* and *P. falciparum* lack the canonical mTOR-based nutrient-sensing pathways present in other eukaryotes but a recent study showed that *P. berghei* is capable of sensing nutrient deprivation by a SNF1-related kinase, KIN1 (Mancio-Silva et al., 2017). *T. gondii* possesses a conserved and uncharacterised homolog of KIN. Determination of protein putatively regulated by KIN in both *Plasmodium* and *T. gondii* may reveal a new set of regulators in adaptation pathways.

Since our observations highlight adaptation of the FASII pathway and the utilization of its acyl-CoA products, it is interesting to note that in plants and bacteria the FASII pathway can be repressed though Acyl-CoA by a feedback loop mechanism (Fujita et al., 2007). Furthermore, supplementation of PUFAs in the fatty acid free diet of mice inhibited both ACCase and FASII (Toussant et al., 1981). Such a mechanism where availability of PUFAs, potentially obtained from the host and its environment, regulates the activity of the FASII would fit with our data. Furthermore, acetylation of FAs has recently been shown to promote the destabilisation of the FA synthesis complex and to inhibit *de novo* FA synthesis and lipogenesis during tumor cell growth (Lin et al., 2016). Acetylome analysis of mutants utilizing acetyl-CoA may reveal how FASII is regulated in nutrient adaptation.

Together, our results reveal the central role of the apicoplast to provide specific precursors for membrane biogenesis during cytokinesis and most importantly to be a central metabolic hub to adapt the parasite metabolic capacities upon nutrient availability and environmental changes. The data also point at major modifications in vesiculation and utilization/scavenging of these membrane structures by the parasite upon such environmental changes. Data also corroborate recent results with the mosquito lipid environment regulates the metabolic activity of transmissible sporozoites (Costa et al., 2018). The fundamental role of these physiological changes induced by the parasite in response to host environment provide novel insights in the parasite biology and offers new avenues to explore in the fight against toxoplasmosis and malaria.

## Ackowledgement

This work and CYB, YYB, SA, NJK, CB are supported by Agence Nationale de la Recherche, France (Grant ANR-12-PDOC-0028-Project Apicolipid), the Atip-Avenir and Finovi programs (CNRS-INSERM-FinoviAtip-AvenirApicolipid projects), and the Laboratoire d’ Excellence Parafrap, France (grant number ANR-11-LABX-0024). CYB and GIM are supported by the LIA CNRS Program (Apicolipid project). MAH is supported by European Research Council (ERC consolidator grant 614880) and the Laboratoire d’Excellence Parafrap, France (grant number ANR-11-LABX-0024).

## Author contributions

SA and NJK designed and performed experiments, analyzed and interpreted data, and wrote the manuscript. LB performed, analyzed and interpreted data for electron microscopy. SD performed Nile Red/LBPA and related immunofluorescence assays. MJS helped performing and analysing the *P. falciparum* lipid starvation growth assay. CB helped performing E. coli complementation assays. BT performed and analysed *T. gondii* proliferation assays and the related statistical analyses. MAH provided the *T. gondii* CrispR-Cas9 system on *Tg*ATS2 and provided experimental advice and supervision regarding this experiment. GIM supervised the *P. falciparum* lipid starvation growth assay. YYB performed, analysed interpreted and supervised lipidomic analyses data and wrote the manuscript. CYB lead the project, designed and interpreted data and wrote the manuscript.

## Materials and Methods

### T. gondii culture

*T. gondii* parental lines RH TATi1-Δ*Ku80* (56) and RH-Δ*Ku80* (57) and derived transgenic cell lines were grown in confluent human foreskin fibroblasts (HFFs) as described (13). Molecular constructs, transfection protocols, growth assays, and egress/invasion assays are described in details in supplemental methods.

### Immunofluorescence assay and Microscopy

Parasites were infected HFF cells grown on coverslips. Primary antibodies used: Mouse anti-HA antibody (InvivoGen, 1:1000), rabbit anti-ACP (1:2000), rabbit anti-TOM40 (1:3000), polyclonal rabbit anti-IMC1 and anti-MIC4 antibodies (1:1000), rabbit anti-Sumo21 at (1:500) and mouse anti-Sag1 (1:500). Secondary antibodies: anti-mouse Alexa488-546, anti-rabbit Alexa546-(Life Technologies, 1:10000). Cells were observed using 63x objective and fluorescent images were acquired with a fluorescence microscope (Axio Imager 2_apotome; ZEISS).

### Plasmodium falciparum growth assays

*P. falciparum* NF54 wild type parasites and FabI-KO (5). Parasite lines were maintained as previously described (58) at 2% haematocrit in RPMI-HEPES supplemented with AlbuMAX II (Gibco). Intra-erythrocytic growth assays in standard media were performed by monitoring the replication of tightly synchronous parasites (5% sorbitol) over four asexual cycles as previously described (59, 60). Media was replaced daily, sub-culturing were performed every 48 h when required, and parasitemia monitored by Giemsa stained blood smears. Growth assays in lipid-depleted media were performed by synchronizing parasites, before transferring trophozoites to lipid-depleted media as previously reported (2, 19). Briefly, lipid-rich AlbuMAX II was replaced by complementing culture media with an equivalent amount of fatty acid free bovine serum albumin (Sigma), 30 μM palmitic acid (C16:0; Sigma) and 45 μM oleic acid (C18:1; Sigma). All assays were performed in triplicates on different days.

### Activity analysis in LPAAT-deficient E. coli strains

*Escherichia coli* strain deficient in LPAAT/AGPAT activity [MS2-1 Δ*plsC*, Coli Genetic Stock Center #7587, Yale University] (29) was used to confirm LPAAT activity in both *Tg*ATS2 and *Tg*AGPAT (details in Supplemental methods)

### Transmission electron microscopy

Parasites were grown for 24 h in Labteks (Nunk, Thermofisher) before fixation in 0.1 M cacodylate buffer with 2.5% glutaraldehyde for 2 h. Samples were then kept in fixative. at 4°C until further processing. Sample were then post-fixed 1h with 1% osmium tetroxide in cacodylate buffer followed by overnight in 2% uranyl acetate in distilled water. After dehydration in graded series of acetonitrile, samples were progressively impregnated in Epon812, the wells were then filled with fresh resin and allowed to polymerize 48 h at 60°C. Ultrathin 70 nm section were obtained with a Leica UCT Ultramicrotome and collected on copper grids. Grids were post-stained with uranyl acetate and lead citrate before their observation on a Jeol1200EXII Transmission Electron Microscope. All chemicals were from Electron Micrsoscopy Sciences.

### Lipidomic analysis by GCMS extraction from T. gondii tachyzoites

Lipidomic analysis were performed as previously described (13, 15, 16). See details in Supplemental methods.

### Stable isotope labelling of T. gondii

Stable isotope labelling using U-^13^C-glucose (Cambridge Isotope Laboratories, USA), lipid extraction, and GC-MS analysis was performed as previously described (13, 15). Freshly infected HFF were incubated in glucose-free medium supplemented with 8 mM U-^13^C-glucose. For FBS starvation study, 5% FBS was add to U-^13^C-glucose medium in standard culture conditions and 1% FBS was add to U-^13^C-glucose medium in starvation culture condition. Parasites were harvested 72 h post-infection and metabolites extracted as above.

## Abbreviations

FA,: fatty acid;
FASII,: fatty acid synthesis;
PL,: phospholipid;
PC,: phosphatidylcholine;
PE,: phosphatidylethanolamine;
PS,: phosphatidylserine;
PI,: phosphatidylinositol;
TAG,: triacylglycerol;
PA,: phosphatidic acid;
GPAT,: glycrol-3-phosphate acyltransferase;
LPA,: lysophosphatidic acid;
AGPAT,: acyl-glycerol3-phosphate acyltransferase;
Drp,: dynamin related protein;
PVM,: parasitophrous vacuole membrane;
KI,: knock-in;
KO,: knock-out;
IMC,: inner membrane complex;
IFA,: immunofluorescent assay;
PM,: plasma membrane;
EM,: electron microscopy;
DAG,: diacylglycerol;
CL,: cardiolipin;
FBS,: foetal bovine serum;
HFF,: human foreskin fibroblast;
ACBP,: acyl-CoA binding protein;
gMVB,: giant multi vesicular body;
LBPA,: lysobisphosphatidic acid;
ACP,: acyl carrier protein;
IPP,: isopentenylphosphate

## Supplementary figure legends

**Supplementary information 1: (A)** Molecular construct for 3’-insertional tagging of *Tg*ATS2 with a C-terminal HA tag. Plasmid harbouring crossover homologous region of gene was linearized with *BlpI* enzyme and transfected into tachyzoites (RH-Δ*ku80*), followed by pyrimethamine drug selection and cloned by limiting dilution. Arrow indicated primer set used for clones screen by PCR. **(B)** PCR analysis using primers specific for the *Tg*ATS2 coding region (left), targeting construct integration (right). **(C)** Western Blot analysis, using anti-HA, confirming the presence of *Tg*ATS2-HA as two major bands of ∼42kDa and ∼60kDa, which correspond to the pre-processed (pATS2, i.e. translated form) and mature (mATS2, i.e. apicoplast located) formsTgATS2. RH-ΔKu80 constitutes the negative control and has no band probed by the anti-HA. GRA1 was used as a loading control. **(D)** Scheme of PCR strategies to isolate a stable Δ*Tg*ATS2-KI and Δ*Tg*ATS2-KO. Fragment of 4000 pb (1) is only amplified in Δ*Tg*ATS2-KI mutants **(E)** but not in RH-Δ*ku80.* Whereas the fragment of 2360 pb (2) is only amplified in *Tg*ATS2-HA line but not in RH-Δ*ku80* and Δ*Tg*ATS2-KO **(F). (G)** Microneme secretion assay was conducted by monitoring the secretion of MIC2 inducted by the A23187 compound or by DMSO as a negative induction control, and measured by Western Blot (TOM40 and GRA1 were used as membrane/pellet and soluble/secreted loading controls, respectively). **(H)** Egress assay, as induced by the A23187 compound showed a mild significant egress defect for the Δ*Tg*ATS2 parasite line (n=3). **(I)** Invasion assay determined via the fluorescent Red-Green assay showed no significant difference between Δ*Tg*ATS2 and parental lines (n=3). **(J)** Immunofluorescence assays investigating the structure and morphology of organelles in the Δ*Tg*ATS2 parasite line: Labelling and observation of mitochondrion (TOM40), nucleus (Sumo21), micronemes (MIC2), parasite PM (Sag1) and inner membrane complex (IMC1) showed no obvious defect in the morphology of these organelles. **(K)** Immunoflorescence assay of the apicoplast (CPN60) showed that Δ*Tg*ATS2 mutants were sometimes lacking their apicoplast or that the organelle was losing integrity (white arrows). Sometimes parasites were attached within a vacuole with one bearing two apicoplasts whilst the other had none (black arrows). Statistical analysis confirmed a statistically significant loss of apicoplast or its compromised integrity between the parental lines and the Δ*Tg*ATS2 mutants (n=3). Scale bars=2 μm. Statistical analyses (Ttest) with *p values* of ≤ 0.05 (*) were considered significant. **(L)** *Tg*AGPAT-HA construct for localization experiment. **(M)** Scheme for CRISPR-Cas9 strategy to disrupt *Tg*AGPAT ORF. Disruption of both *Tg*ATS2 and TgAGPAT using CRISPR-Cas9 affect apicoplast biogenesis. Immunofluorescence assay performed 48 h post-transfection of RH-Δku80 **(N)** and ΔTgATS2 parasites (O) with pTOXO-Cas9-GFP-CRISPR∷gTgAGPAT, using anti-CPN60 antibody as apicoplast marker. Scale bars=2 μm. Yellow arrows indicate a correct apicoplast segregation in tachyzoites and white arrows indicate apicoplast structure loss.

**Supplementary information 2:** Fatty acid composition of PC **(A)**, PI **(B)**, PE **(C)** PG **(D)**, CL **(E)**, and PS **(F)** molecular species in Δ*Tg*ATS2 (red) and parental line (blue). Fatty acid composition of HFF cells **(G)** and FBS **(H). (I)** Plaque assay performed with the parental line and Δ*Tg*ATS2 complemented with 20 μM of PA(14:0/14:0) or PA(16:0/18:1) suggests a increase in parasite proliferation in the parental line complemented with both PA sources but not in the Δ*Tg*ATS2 mutant line. **(J)** Growth fitness analysis performed using semi-automated cell-based assay on infected cells with parental line and Δ*Tg*ATS2 30 h post-infection confirmed a statistical increase in the intracellular development of the PA-complemented parental line but not in Δ*Tg*ATS2. Parasite numbers within vacuoles were counted to obtain the proliferative index, which was evaluated by parasite/vacuole number ratio, in two independent Δ*Tg*ATS2 lines conducted in triplicate. *, *p* ≤ 0.05 were considered significant.

Fluorescent lipid import assay on extracellular parental (*Tg*HX) and Δ*Tg*ATS2 parasites supplied with either PA(C18:1/C12:0-NBD) or PC(C18:1/C12:0-NBD) as a control. The incorporation of fluorescent NBD-PA **(K)** or NBD-PC **(L)** in the parasite membrane was quantified by measuring fluorescence intensity. Δ*Tg*ATS2 imported significantly more PA and PC than the parental control line. Fluorescent PA import was higher than the PC import in the Δ*Tg*ATS2 mutant **(M)**.

Supplementary information 3: Multiple sequence alignment and protein sequence analysis of *Tg*DrpC

**(A)** *Tg*DrpC is the closes homolog of the human mitochondrial Dynamin-like 1 (*Hs*Drp1). HsDrp1 function and regulation depends on its interaction with PA within its Stalk domain (blue insert), and more particularly a variable domain (dark blue insert) as well as hydrophobic residues in its insertional loop/PAbinding domain (black insert). *Tg*DrpC posseses a conserver Stalk and hydrophobic PA binding domain, indicating its is capable of interacting with PA similarly as HsDrp1. None of *Tg*DrpA, and *Tg*DrpB has a similar conserved variable and PA binding domains **(B, C)**.

**Supplementary information 3:** Scheme depicting the molecular construct for the endogenous HA tagging of *Tg*DrpC using CrispR-Cas9 approach **(A)** and endogenous GFP tagging of TgDrpA using CrispR_Cas9 approach **(B).** *Tg*ATS2 disruption does not affect *Tg*DrpA localisation **(C)** Endogenous expression of a TgDrpA-GFP sequence shows the typical localisation of the protein at the apicoplast of tachzoite in the parental (wt) line. Similar expression of TgDrpA in the Δ*Tg*ATS2 mutant background did not lead to any mislocalisation of the apicoplast-resident protein.

**Supplementary information 4:** Mass isotopolog distribution of ^13^C into C14:0 (A), C16:0 (B) and C18:0 (C) after U-^13^C-glucose labelling in FBS starvation (0.2% FBS final concentration). Experiments were conducted in triplicates. * *P values* of ≤ 0.05 from statistical analyses (Ttests) were considered statistically significant. (D) Δ*Tg*ATS2 intracellular parasite seem not to be affected by FBS starvation. Immunofluorescence assay on parental line and Δ*Tg*ATS2 grown 24 h in FBS starvation conditions 0% and 0.25%, using anti-IMC1/Alexa546 and anti-Sag1/Alexa488 antibodies. Scale bars=2 μm

**Supplementary information 5:** Scheme depicting the molecular construct to generate tagged inducible knock-downs of *Tg*ACBP1 (**A**) and *Tg*ACBP2 (**B**) (red arrows and their respective numbers represent the primers used to probe the constructs and parental lines). *Tg*ACBP1-iKO and *Tg*ACBP2-iKO clones were confirmed by PCR (C,D) using the primers as indicated **E,F**. Expression of endogenously tagged *Tg*ACBP1-HA and *Tg*ACBP2-GFP determine their respective localisation at the cytosol for *Tg*ACBP1 **(G)** and at the mitochondrion for *Tg*ACBP2 **(H)** Western Blots confirm the down-regulation of the inducible knock-downs of *Tg*ACBP1-HA **(I)** and *Tg*ACBP2-GFP **(J)** by ATc treatment.

## Supplemental methods

### Gene identification and sequence analysis

*Arabidopsis thaliana* sequence of ATS2 (GenBank™ and TAIR™ IDs: NP_194787 and AT4G30580 respectively) was used as a query sequences for BLAST searches against the *Toxoplasma gondii* genome on ToxoDB database (www.toxodb.org). Phylogenetic analysis of AGPAT related proteins was performed on the Phylogeny.fr platform (1) using proteins described in **Table1**. Protein sequences were then aligned by ClustalW software (2) and the maximum likelihood phylogeny was generated using the PhyML (3). We generated multiple sequence alignment using Clustal Omega (4).

### Toxoplasma gondii plasmid constructs

Plasmid LIC-3HA-DHFR was used to generate a 3’ endogenous tagging with 3xHA coding sequence of TGME49_297640 (*TgATS2*) and TGME49_240860 (*TgAGPAT*). A 2229 bp fragment corresponding to the 3’ of *Tg*ATS2 was amplified from genomic DNA using primer sets 5’-TCCTCCACTTCCAATTTTAGCGTTCGTCTCGGTGGCGGC-3’ and 5’-TACTTCCAATCCAATGCTTCAGACACTCGGTGCAAA-3. A 5466 bp fragment corresponding to promoter and gene sequence of *Tg*AGPAT was amplified using primer sets 5’-TACTTCCAATCCAATGCAGCCAGCAAAGGACGAAAGG-3’ and 5’-TCCTCCACTTCCAATTTTAGCGAGACCGTGGCCTCGGTGGG-3’. These fragments were cloned into pLIC-3HA-DHFR vector as described previously [Huynh and Carruthers 2009]. Vectors LIC-*Tg*ATS2-3HA-DHFR and LIC-*Tg*AGPAT-3HA-DHFR were confirmed by PCR screen using primer sets 5’-GCATAATCGGGCACATCATA-3’ and 5’-ATACGCATAATCGGGCACATCATA-3’and by sequencing (GATC Biotech™).

Plasmid pTOXO_Cas9-CRISPR (gift from Hakimi Lab, Grenoble, France) was used to integrate a gRNA within *BsaI* restriction site as previously described (5). Briefly, Crisp-Fwd and Crisp-Rv primer sets were phosphorylated and annealed: *TgATS2-KI*: 5’-AAGTTACGGGTGTGCGCCGCCTTGCG-3’ and 5’-AAAACGCAAGGCGGCGCACACCCGTA-3’, *TgATS2-KO*: 5’-AAGTTGGAGCGCCGACGGGCGACTGG-3’ and 5’-AAAACCAGTCGCCCGTCGGCGCTCCA-3’, *TgAGPAT-KO*: 5’-AAGTTCTCTGCCGAGTTCCAATCGCG-3’ and 5’-AAAACGCGATTGGAACTCGGCAGAGA-3’. The gRNAs were then ligated into pTOXO_Cas9-CRISPR plasmid linearized with *BsaI*, yielding pTOXO_Cas9-CRISPR∷g*Tg*ATS2-KI, pTOXO_Cas9-CRISPR∷g*Tg*ATS2-KO and pTOXO_Cas9-CRISPR∷g*Tg*AGPAT-KO, respectively.

For *Tg*ACBP1iHA KD, the 5’ UTR flank was PCR amplified using primers below and inserted into ApaI/NdeI sites of pPR2-HA3 (6). The 3’ flank was then inserted in frame with a Tet7O/SAG4 promoter using XmaI/NotI sites. Plasmid was linearized with NotI prior to transfection. For ACBP2, flanks were PCR amplified using respective primers below into the plasmid pPR2-GFP) or pPR2-mCherry (adapted from pPR2-HA3, (6). The 5’ UTR flank was inserted first using ApaI/NdeI sites. Next, the 3’ UTR flank was inserted using MscI/NotI sites. ACBP2 cDNA sequence was inserted last into BglII/AvrII sites. ACBP1 pLIC was PCR amplified using pLIC primers into pLIC-HA3-CAT (7), and linearized prior to transfection and selection on chloramphenicol. DrpA and DrpC were localized by CRISPR Cas9 strategy. Guides were inserted into Cas9 U6 universal plasmid (5) by either standard ligation of annealed primers or Q5 mutagenesis. Cells were transfected together with PCR product encoding either HA3-CAT and DrpC homology flanks, for DrpC or GFP sequence without selection and DrpA homology flanks for DrpA.

For DrpC HA3 CAT CRISPR Cas9 tagging, DrpC was tagged at the 3’ terminus by CRISPR Cas9 (5). Primers FOR and REV were annealed together and ligated into U6 universal plasmid (5). The HA3-CAT cassette was PCR amplified by primers with 50 bp homology flanks corresponding to the 3’ end of DrpC and in frame with HA3. 50 μg of both plasmid and PCR product were transfected and placed under chloramphenicol selection (8). DrpC was also localized by pLIC-HA3-CAT (7) using For *Tg*ACBP1 iHA KD strain, the 5’ UTR flank was PCR amplified using primers below and inserted into ApaI/NdeI sites of pPR2-HA3 (6). The 3’ flank was then inserted in frame with a Tet7O/SAG4 promoter using XmaI/NotI sites. Plasmid was linearized with NotI prior to transfection.

For *Tg*ACBP2 iKD GFP strain, flanking regions were PCR amplified using respective primers below into the plasmid pPR2-GFP) or pPR2-mCherry (adapted from pPR2-HA3, (6)Katris et al. 2014. The 5’ UTR flank was inserted first using ApaI/NdeI sites. Next, the 3’ UTR flank was inserted using MscI/NotI sites. ACBP2 cDNA sequence was inserted last into BglII/AvrII sites.

For *Tg*ATS2 knockout by CRISPR-CAs9, an appropriate HXGPRT cassette amplified by PCR from pMini (kind gift from the Hakimi laboratory) using those primer sets, *TgATS2-KI*: 5’-GAGGCCCTGCGTCTCCTCAAGCGAAAGGCGCCGCCACAGTCGACGGGTGTGC GCC-GCCTCAGCACGAAACCTTGCATTCAAACC-3’ and 5’-GCTACTCCTTCTTCCCCTCTCG-CGTTGTGTGTCTCCCCGTCGCGTTCTGCGTCGCCAGCAGTGTCACTGTAGCCTGCCAGAACA-3’; *TgATS2-KO*: 5’-GACACACAACGCGAGAGGGGAAGAAGGAGTAGCTCTCG-TCGCCTTTCCAGAAGGTACTCCAGCACGAAACCTTGCATTCAAACC-3’ and 5’-CTTCG-CTGCTCGTTCGTCTTCATGTGGGGAAGGAGCAGCACGAAACCTTG-CATTCAAACC-3’.

### Toxoplasma gondii transfection

RH-Δ*Ku80* parasite line was transfected with 100 μg of pLIC-*Tg*ATS2-3HA-DHFR linearized with *BlpI* for stable integration of HA-tag at C-terminus of *Tg*ATS2. 150 μg pTOXO_Cas9-CRISPR∷g*Tg*ATS2-KO and pTOXO_Cas9-CRISPR∷g*Tg*AGPAT-KO were transfected in *Tg*ATS2-HA line with 10 μg of appropriate HXGPRT cassette for TgATS2-KI and TGATS2-KO, PCR product as described above. Electroporations were performed in a 2-mm cuvette in a BTX ECM 630 (Harvard Apparatus, at 1,100 V, 25 Ω, and 25 μF. Stable lines expressing the tagged constructs were selected selected in media with 1 μM pyrimethamine or 25 μg/ml mycophenolic acid and 50 μg/ml xanthine and cloned by limiting dilution.

RH-Δ*Ku80* parasites were also transiently transfected with pLIC-*Tg*AGPAT-3HA-DHFR. pTOXO_Cas9-CRISPR∷g*Tg*AGPAT-KO was transfected in RH-Δ*Ku80* parasites for a simple mutant Δ*TgAGPAT* and in Δ*TgATS2 parasites* to obtain a double mutant Δ*TgATS2/*Δ*TgAGPAT*. The plasmid pMORN1-CherryRFP-MORN1/SagCAT were transfected in both RH-Δ*Ku80* and Δ*TgATS2* parasite lines. All transfections were performed with 150 μg of DNA and electroporation conditions were as described above. Transfected parasites were incubated at different concentration with HFF cell 48 h prior to immunofluorescence assay.

### T. gondii growth assays

*-Plaque Assay:* HFF monolayers were infected with 500 parasites and allowed to develop for 10 days before staining with Crystal Violet (Sigma) and cell growth assessment by light microscopy for the presence of intact HFF.

*-Cell-based assay: T. gondii* growth was determined with an automatic microscope-based screening (Olympus ScanR, Japan). HFFs were seeded at a density of 10,000 cells per well into 96-well plates and were allowed to grow and equilibrate for 48 h at 37°C. Cells were then infected with 4 × 10^4^ parasites/well. Invasion was synchronized by briefly centrifugation of plate at 250 *g* and placed at 37°C for 2 h. The assay was run for 30 h. Hoechst 33342 (Life technologies) stain was then loaded on live cells/parasites at 5 μg/ml for 20 min. Infected cells were fixed with PFA (3.7%) for 10 min at 37°C. A mouse anti-GRA1/Alexa488 labeling (dilution 1:500) was used to identify parasitophorous vacuoles. A total of 20 fields per well were taken using the 20X objective. Images were collected for the distinct fluorescence channels (Hoechst 33342: eg. 360-370 nm, em. 420-460 nm and Alexa488: ex. 460-495, em. 510-550 nm). Images were then analyzed using the ScanR analysis software (Olympus, Tokyo, Japan). For Alexa488 channels images (vacuoles) an intensity algorithm module was used where a fixe threshold was defined with a minimum of 100 pixels size in order to segment the smallest vacuoles (one or two parasite). For Hoechst channel images (parasites nuclei), image process consists to apply a strong background correction and detected parasites with an edge algorithm. A minimum object size of 5 pixels and a maximum object 20 pixel larger one was chosen to discriminate each parasite. ScanR analysis module interface as in flow cytometry allow us to extract and display data as scatter plots and histograms. Using a “gating” procedure we were able to hierarchically filter selected data points with precise boundaries (e.g. number of vacuoles vs number of parasite/vacuoles). The proliferative index was evaluated by parasite/vacuole number ratio.

### Activity analysis in LPAAT-deficient E. coli strains

Coding sequence of *TgATS2* was synthesized (Genscript). *TgAGPAT* coding sequence was amplified by RT-PCR using primer sets 5’-ATGGCGTCCACGCCGCTGC-3’/5’-TTAGAGACCGTGGCCTCGGTG-3’ and *TgAGPAT*_Δ*N-ter1-72*_ coding was amplified by RT-PCR using primer sets 5’-CTCAACCGCCCGCCCAGGAATTA-3’/5’-TTAGAGACCGTGGCCTCGGTG-3’.

These sequences were digested and ligated into *HindIII* restriction site on pQE30Xa vector (Quiagen) to generate expression vectors. Additionally, gene coding for *E. coli* LPAAT activity *plsC*, was amplified from *E. coli* DH5alpha genomic DNA using primer sets 5’-CTATATATCTTTCGTCTTATTATTAC-3’/ 5’-AACTTTTCCGGCGGCTTC-3’ and ligated into pQE30Xa vector. Then these acyltransferase vectors and empty pQE30Xa vector as negative control were transfected to eletrocompletent cells of MS2-1 Δ*plsC* deficient *E.coli.* pREP4 repressor vector to regulate Lac promotor activity. Transformed bacterial populations were grown at 37°C in order to promote growth of all isolates. Two independent clones of each bacterial strain that harbours each plasmid-of-interest were isolated for this study. Rescue of LPAAT activity in SM2-1Δ*plsC* mutant was measured by the ability to grow at elevated temperature, 42°C, non-permissive temperature in LB medium as previously described (9). Bacteria were first grown in LB media at 37°C to stationary phase, then the cultures were diluted to OD_600_=0.04 and finally inoculated with several dilutions (at 10^−1^ to 10^−6^) on LB plates and incubated for 24 h at permissive (30°C) and non-permissive (42°C) temperatures. All experiments were conducted in triplicate with both independent clones.

### Parasite egress assay

WT or Δ*Tg*ATS2 parasites were incubated on HFF cells for approximately 26 h before aspirating medium and replacing with DMEM containing 2 μM A23187 or DMSO. Parasites were incubated for 3 min before addition of an equivalent volume of 2x fixative containing 5% Paraformaldehyde, 0.05% glutaraldehyde in PBS (final concentration 2.5% Paraformaldehyde, 0.025% glutaraldehyde). Cells were fixed for 15 minutes before permeabilizing with 0.025% TX100 in PBS for 10 min and then Blocking overnight in blocking solution (2% FBS in PBS). Samples were then probed by immunofluorescence assay and counted manually for egress.

### Red/Green parasite invasion assay

Experiment was performed as per (6). Parasites were grown for 2 days and harvested intracellular after replacing medium with ENDO buffer (44.7 mM K_2_SO_4_, 10 mM MgSO4, 106 mM sucrose, 5 mM glucose, 20 mM Tris-H_2_SO_4_, 3.5 mg/ml BSA, pH 8.2). Cells were scraped, needle passed, filtered and centrifuged at 1800 rpm for 10 min. Cells were resuspended to a concentration of 2.5 × 10^7^ cells ml^−1^ in ENDO buffer and settled for 20 min onto host cells. Once settled, medium was aspirated and replaced with Invasion buffer (DMEM, 3% FBS and 10 mM HEPES). Parasites were allowed to invade for 15 min before fixation with 2.5% Paraformaldehyde and 0.02% glutaraldehyde. Samples were then blocked in 2% FBS in PBS overnight at 4°C. Samples were probed with mouse anti-SAG1, before washing with PBS, then permeabilized with 0.25% Triton-X100 in PBS. Cells were then probed with rabbit anti-GAP45 and washed in PBS. Samples were then probed with Alexafluor anti-mouse 546 and anti-rabbit 488 before mounting onto slides. Cells were imaged by microscopy and invasion rate determined using ImageJ.

### Lipidomic analysis by GC-MS

Lipid extraction and analysis of tachyzoites was performed as previously described (10-12). Freshly egressed tachyzoites (1 × 10^8^ cell equivalents) grown in standard culture or in starvation culture, were metabolically quenched by rapid chilling of the cell suspension in a dry ice/ethanol bath and lipids were extracted in chloroform/methanol/water (2:1:0.8, v/v/v containing 25 nmol tridecanoic acid C13:0 as extraction internal standard) for total lipid analysis.

*-For lipid quantification:* Total lipid extraction was performed as described previously (11). Parasites were prepared as described above except for the addition of 0.1 M of HCl to promote PA and LPA extraction. Pooled organic phase was subjected to biphasic separation by adding 0.1 M HCl. In both protocols, the organic phase was dried with speed vaccum and dissolved in 1-butanol.

*-Total lipids analysis:* An aliquot of the lipid extract was dried in vacuum concentrator with 1 nmol pentadecanoic acid C15:0 as internal standard. Then the dried lipid was dissolved in the chloroform/methanol, (2:1, v/v) and derivatised with MethPrep II (Alltech). The resulting fatty acid methyl esters was analysed by GC-MS as described previously (11). Fatty acid methyl esters were identified by their mass spectrum and retention time compared to authentic standards.

*-Lipid quantification:* Total lipid fraction was separated by 2D-HPTLC with 5 μg PA(C17:0/C17:0) and 5 μg LPA(C17:0) (Avanti Polar lipids) using chloroform/methanol/28% NH4OH, 60:35:8 (v/v) as the 1st dimension solvent system and chloroform/acetone/methanol/acetic acid/water, 50:20:10:13:5 (v/v/v/v/v) as the 2nd dimension solvent system (11). For DAG analysis, total lipid fraction was separated by 1D-HPTLC using hexan/diethlether/formic acid, 80:20:2 (v/v/v) as solvent system. The spot on the HPTLC corresponding to each lipid was scrapped off and lipids were directly derivatised with 0.5 M methanoic HCl in the presence of 1 nmol pentadecanoic acid (C15:0) as internal standard. The resulting fatty acid methyl esters were extracted with hexane and analysed by GC-MS (11). Resulted FAME and cholesterol-TMS was analyzed by GC-MS (5977A-7890B, Agilent). FAME was then quantified using Mass Hunter Quantification software (Agilent). All statistical analyses were conducted using GraphPad Prism software. *P values* of ≤ 0.05 from statistical analyses (Ttests) were considered statistically significant.

### Phospholipid import assay

Freshly lysed cultures of WT or Δ*Tg*ATS2 parasites were harvested, filtered and resuspended in DMEM to a concentration of approximately 2 × 10^8^ cells ml^−1^. Cells were then mixed with a 2x solution containing 10μg ml^−1^ NBD-PA or NBD-PC (5μg ml^−1^ final) and incubated at 37°C. Parasites were then spun down, resuspended in PBS. PFA was then added to a final concentration of 2.5%, and cells were fixed for 15 min before being spun down again and resuspended in 1xPBS. Parasites were smeared onto polyethyleneimine (PEI) coated coverslips (6), and then probed with anti-SAG1 primary and Alexfluor anti-mouse 546 secondary antibodies by immunofluorescence microscopy, stained with DAPI and mounted onto slides. Samples were imaged by microscopy. SAG1 labelling was used to identify parasites using ImageJ and then estimate the amount of NBD-lipid uptaken by the parasites.

