## Supplemental figures for "Division and adaptation to host nutritional environment of apicomplexan parasites depend on apicoplast lipid metabolic plasticity and host organelles remodelling"

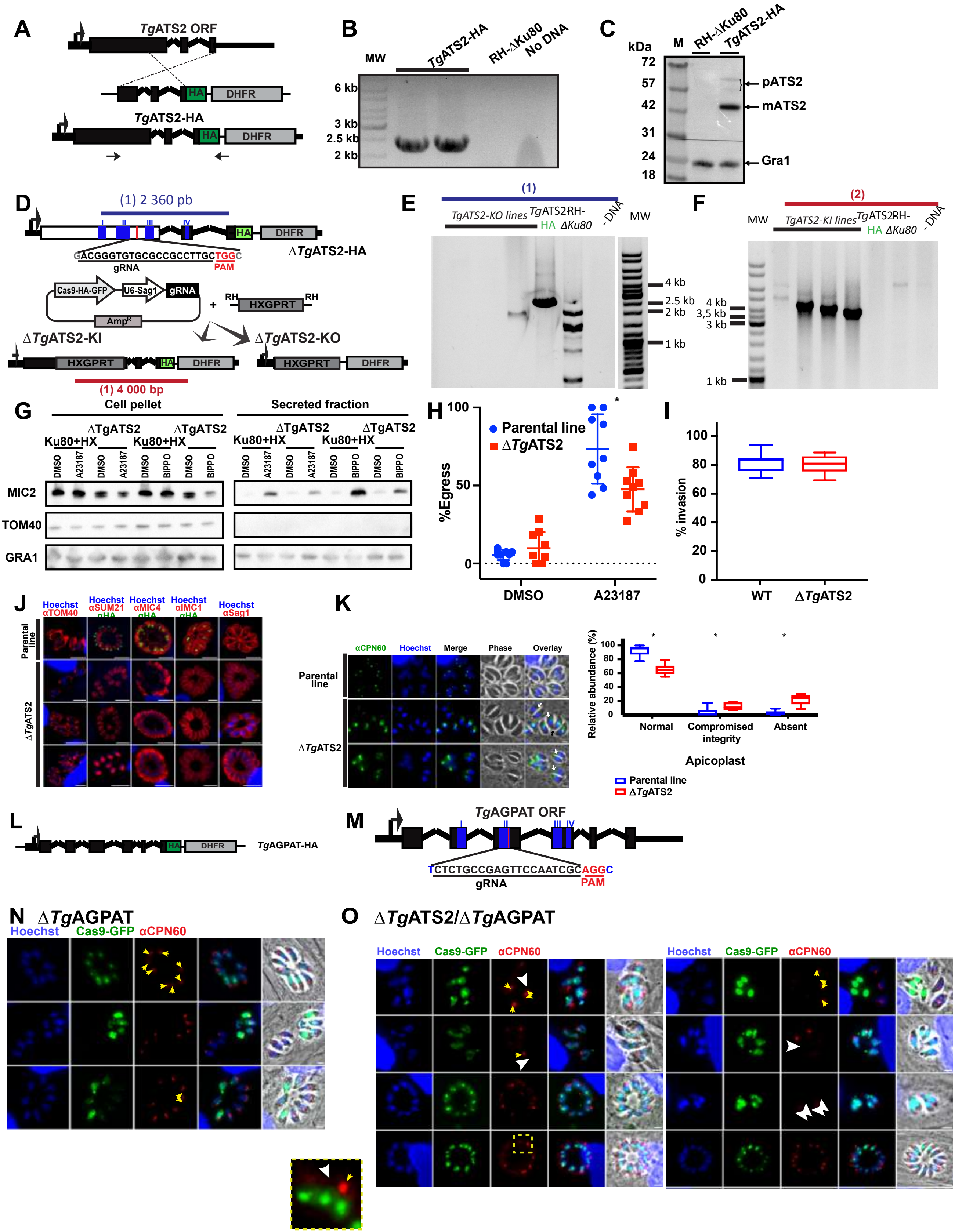
A
TgATS2 ORF
HA DHFR
TgATS2-HA
B
MW
TgATS2-HA
RH-ΔKu80
No DNA
6 kb
3 kb
2.5 kb
2 kb
C
kDa
M
RH-ΔKu80
TgATS2-HA
72
57
42
31
24
18
pATS2
mATS2
Gra1
D
(1) 2 360 pb
HA DHFR
GACGGGTGTGCGCCGCCTTGCTGGC
gRNA
PAM
ΔTgATS2-HA
Cas9-HA-GFP
U6-Sag1
gRNA
AmpR
RH HXGPRT RH
ΔTgATS2-KI
ΔTgATS2-KO
HXGPRT HA DHFR
HXGPRT DHFR
(1) 4 000 bp
E
(1)
TgATS2-KO lines
TgATS2RH-HA ΔKu80
- DNA
MW
4 kb
2.5 kb
2 kb
1 kb
F
(2)
MW
TgATS2-KI lines
TgATS2RH-HA ΔKu80
- DNA
4 kb
3,5 kb
3 kb
1 kb
G
Cell pellet
Secreted fraction
Ku80+HX
ΔTgATS2
DMSO
A23187
BIPPO
MIC2
TOM40
GRA1
H
Parental line
ΔTgATS2
%Egress
100
50
0
DMSO
A23187
I
% invasion
100
80
60
40
20
0
WT
ΔTgATS2
J
Hoechst αTOM40
Hoechst αSUM21 αHA
Hoechst αMIC4 αHA
Hoechst αIMC1 αHA
Hoechst αSag1
Parental line
ΔTgATS2
K
αCPN60
Hoechst
Merge
Phase
Overlay
Parental line
ΔTgATS2
Relative abundance (%)
100
80
60
40
20
0
Normal
Compromised integrity
Absent
Apicoplast
Parental line
ΔTgATS2
L
HA DHFR
TgAGPAT-HA
M
TgAGPAT ORF
TCTCTGCCGAGTTCCAATCGCAGGC
gRNA
PAM
N
ΔTgAGPAT
Hoechst
Cas9-GFP
αCPN60
O
ΔTgATS2/ΔTgAGPAT
Hoechst
Cas9-GFP
αCPN60

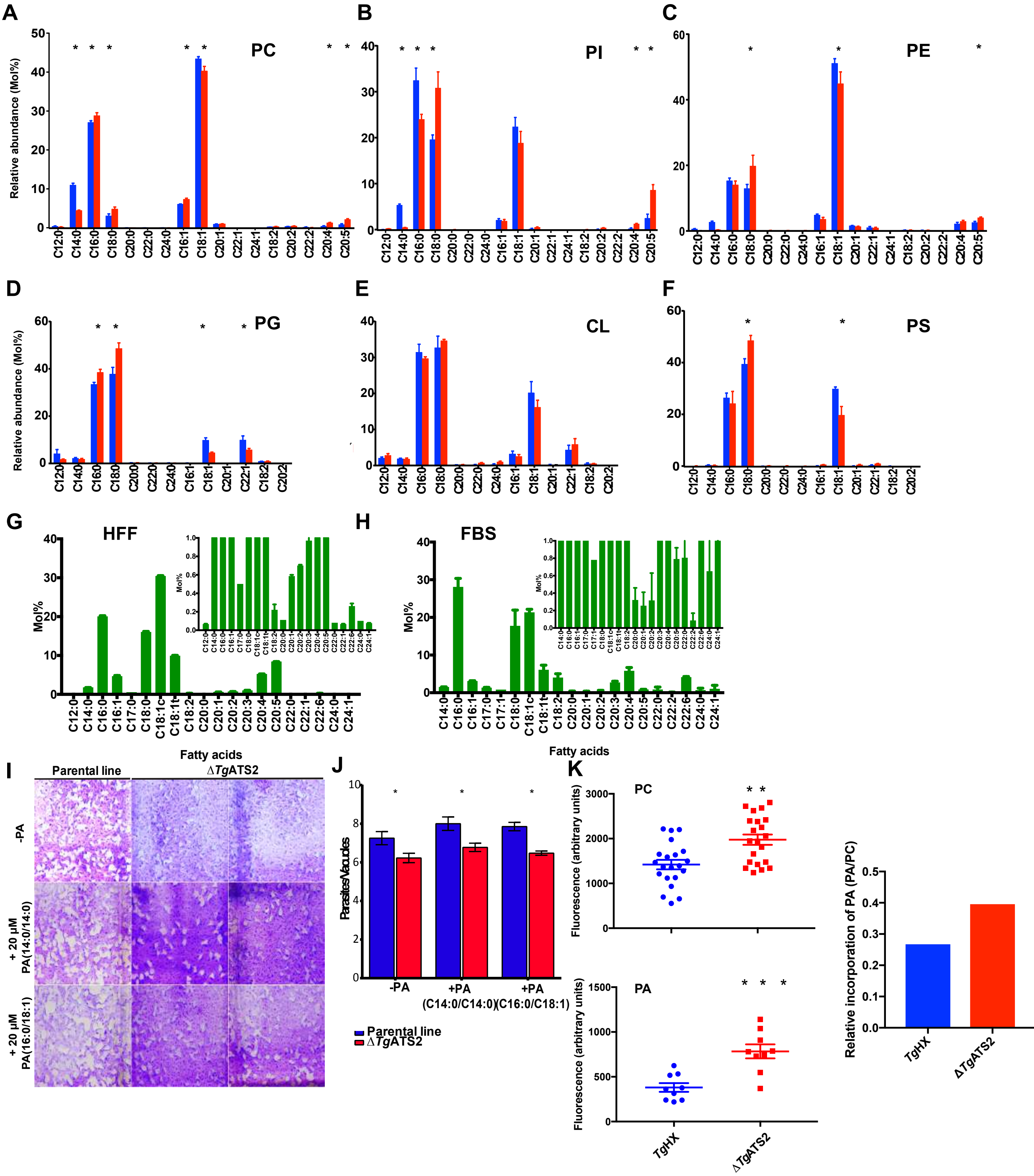

A
PC
Relative abundance (Mol%)
B
PI
C
PE
D
PG
E
CL
F
PS
G
HFF
Mol%
H
FBS
I
Parental line
Fatty acids
ΔTgATS2
-PA
+ 20 µM PA(14:0/14:0)
+ 20 µM PA(16:0/18:1)
J
-PA
+PA (C14:0/C14:0)
+PA (C16:0/C18:1)
Parental line
ΔTgATS2
Fatty acids
K
PC
PA
Fluorescence (arbitrary units)
TgHX
ΔTgATS2
Relative incorporation of PA (PA/PC)

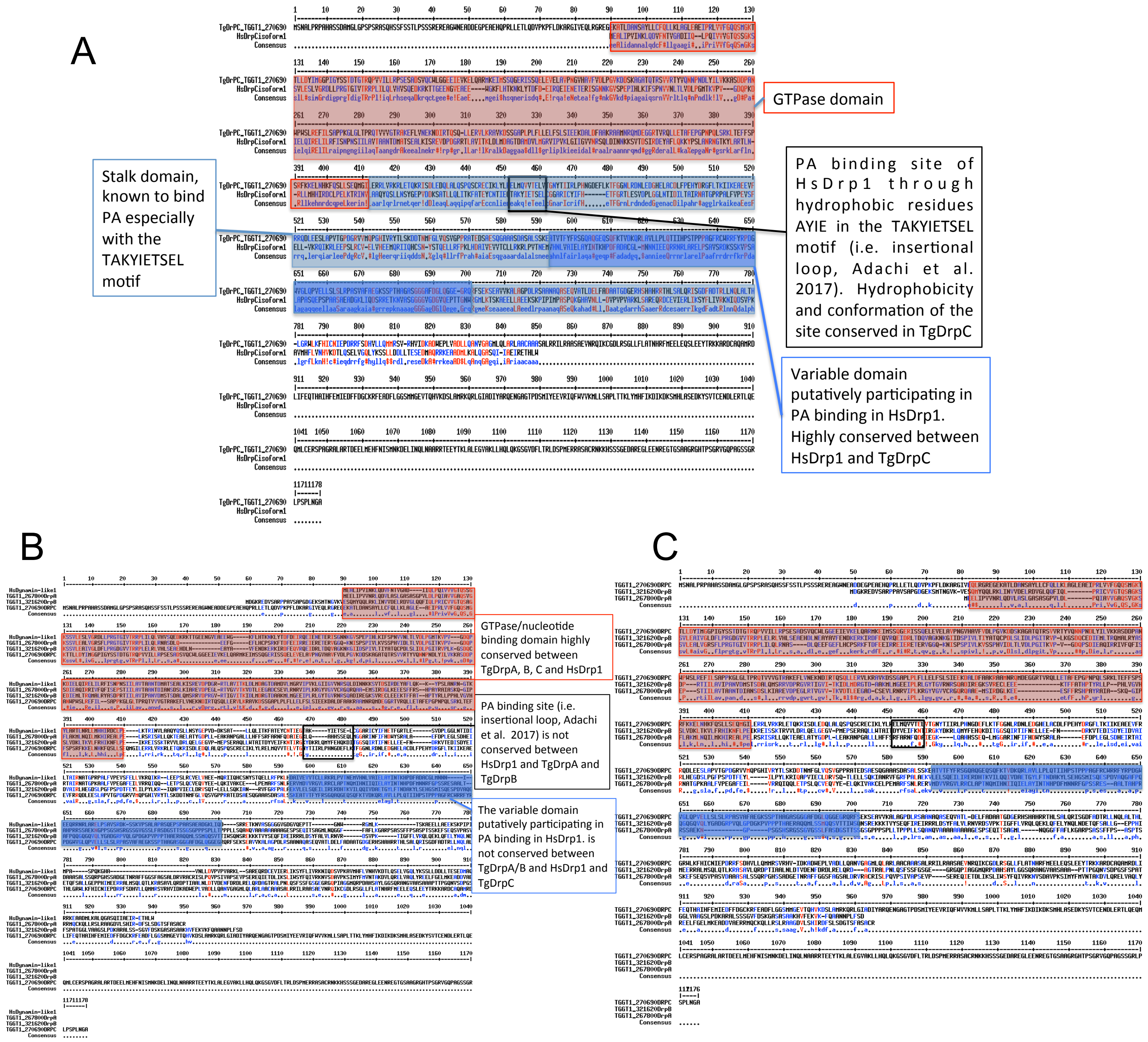

A
GTPase domain
Stalk domain, known to bind PA especially with the TAKYIETSEL motif
PA binding site of HsDrp1 through hydrophobic residues AYIE in the TAKYIETSEL motif (i.e. insertional loop, Adachi et al. 2017). Hydrophobicity and conformation of the site conserved in TgDrpC
Variable domain putatively participating in PA binding in HsDrp1. Highly conserved between HsDrp1 and TgDrpC
B
GTPase/nucleotide binding domain highly conserved between TgDrpA, B, C and HsDrp1
PA binding site (i.e. insertional loop, Adachi et al. 2017) is not conserved between HsDrp1 and TgDrpA and TgDrpB
The variable domain putatively participating in PA binding in HsDrp1. is not conserved between TgDrpA/B and HsDrp1 and TgDrpC
C

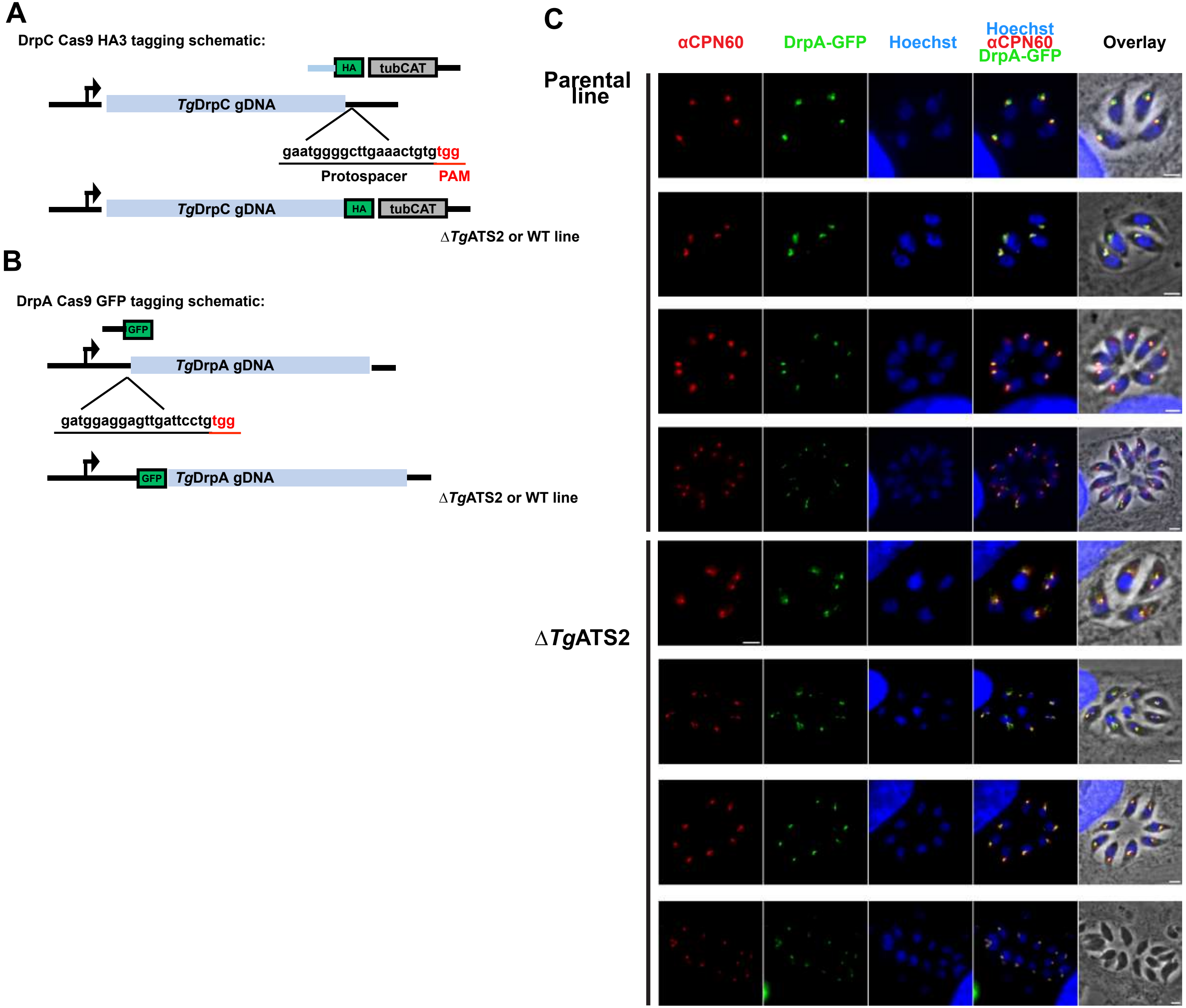
A
DrpC Cas9 HA3 tagging schematic:
HA
tubCAT
TgDrpC gDNA
gaatggggcttgaaactgtgtgg
Protospacer
PAM
TgDrpC gDNA
HA
tubCAT
ΔTgATS2 or WT line
B
DrpA Cas9 GFP tagging schematic:
GFP
TgDrpA gDNA
gatggaggagttgattcctgtgg
GFP
TgDrpA gDNA
ΔTgATS2 or WT line
C
αCPN60
DrpA-GFP
Hoechst
Hoechst
αCPN60
DrpA-GFP
Overlay
Parental line
ΔTgATS2

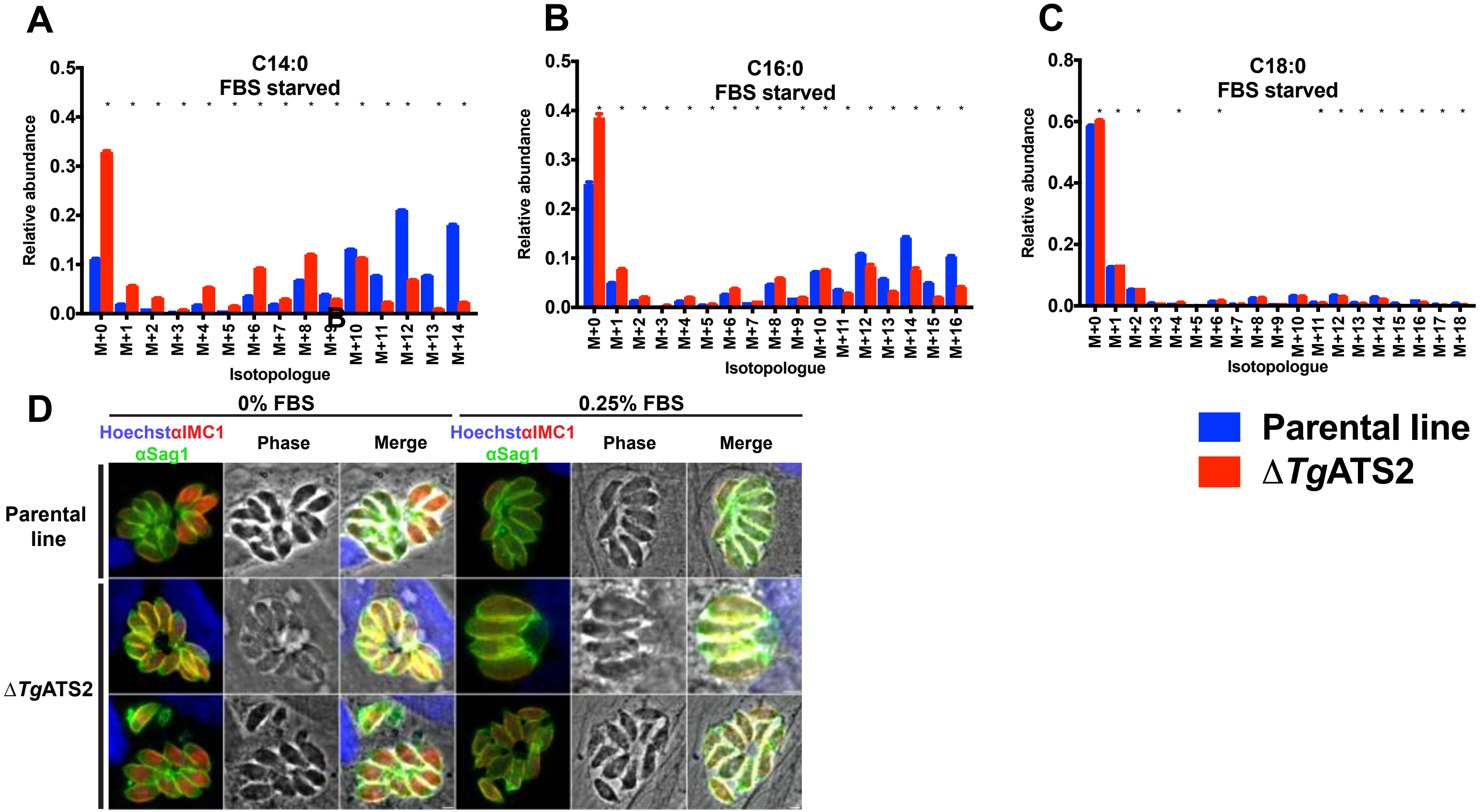
A
C14:0
FBS starved
Relative abundance
Isotopologue
B
C16:0
FBS starved
C
C18:0
FBS starved
D
0% FBS
0.25% FBS
Hoechst αIMC1 αSag1
Phase
Merge
Parental line
ΔTgATS2
Parental line
ΔTgATS2

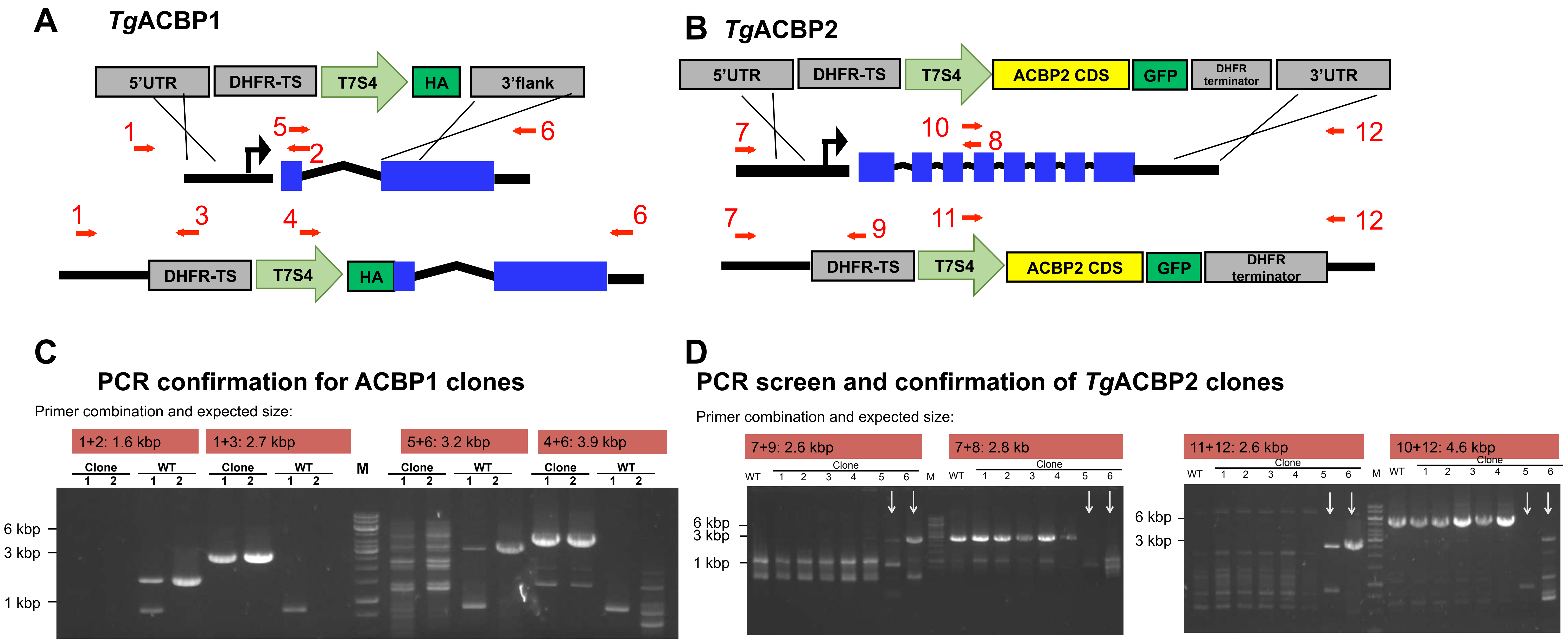


**E**

| ACBP1 primer number | Primer sequence |
|---|---|
| #1 | TTGACTCGTCTCCACATGAG |
| #2 | GGTGGCTGGATAGTTGG |
| #3 | TCTTCTTTGAGGGAAGAGG |
| #4 | GGTACCGAGCTCGACTTTCAC |
| #5 | CCCAACTATCCAGCCACC |
| #6 | GCATGTCGAAGGAGCAGTC |

**F**

| ACBP2 primer number | Primer sequence |
|---|---|
| #7 | ACGGAGAGCGTTTGAGAGG |
| #8 | CGAACACATACAGGAACTTCAGG |
| #9 | TGTCTTCTTCTTTGAGGGAAGAGG |
| #10 | GAAGTTCCTGTATGTGTTCGTGC |
| #11 | GGTACCGAGCTCGACTTTCAC |
| #12 | CAACCGTACTTTCATCCTTGG |

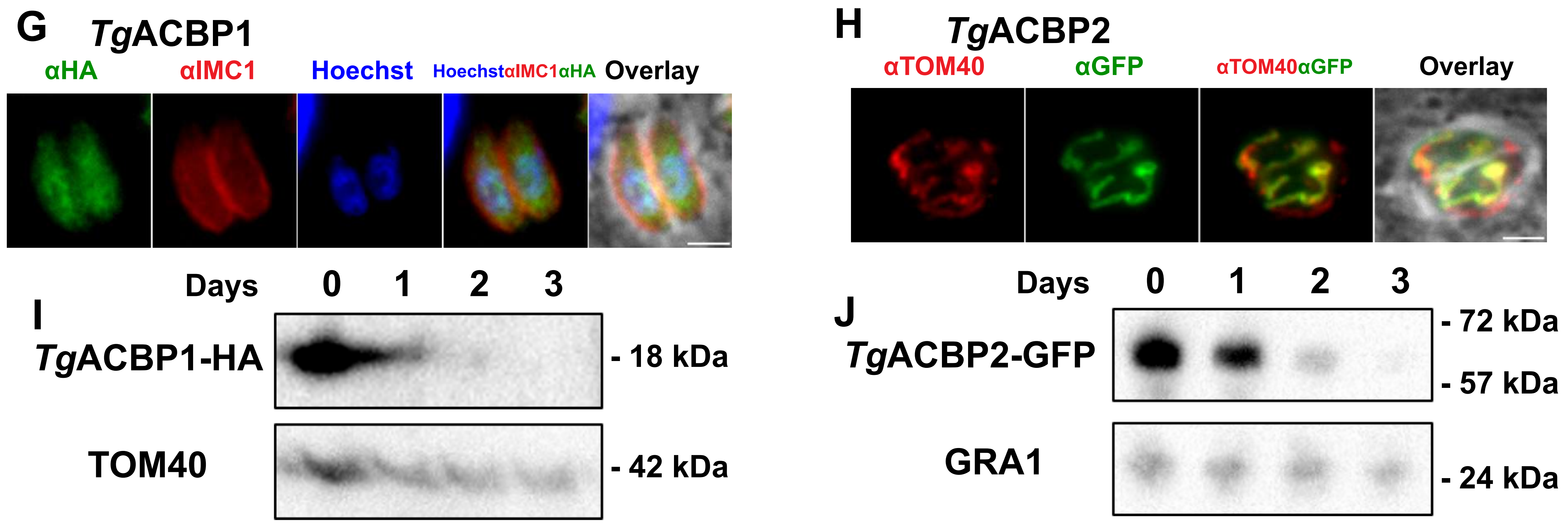

| Enzyme name | | Pathway | *T. gondii* Accession numbers (http://toxodb.org/toxo/) | *T. gondii* Phenotype score in10% FBS (Siddick et al. 2016) | *P. falciparum* Orthologue accession numbers (http://plasmodb.org/plasmo/) | *P. falciparum* Blood stage mutability in rich culture medium (Zhang et al. 2018) | *P. falciparum* Blood stage peak expression profile (PlasmoDB) | *P. falciparum* Blood stage transcriptomic upregulation in starved patients (Daily et al. 2007) | *P. yoelii* Orthologue accession numbers (http://plasmodb.org/plasmo/) | *P. yoelii* Late liver stage expression profile (Tarun et al. 2008) | Reported essentiality: |
|---|---|---|---|---|---|---|---|---|---|---|---|
| oTPT | Outer membrane triose phosphate translocator | precursor transporter | TGGT1_261070 * | -4.44 | PF3D7_0508300 | Non-mutable | Trophozoite (30 h) | (+) 2X | PY00389 | (+);(-); 40 h | * Essential (Brooks et al. 2010) |
| iTPT | inner membrane triose phosphate translocator | precursor transporter | | | PF3D7_0530200 | Mutable | Trophozoite (32 h) | (++) 8X | PY01812 | (+);(+); 40 h | |
| Pyruvate kinase | pyruvate kinase | generation of pyruvate | TGGT1_299070 | -2.76 | PF3D7_1037100 | Non-mutable | Trophozoite (24 h) | (++) 6.5X | PY03879 | (+);(+); 40-50 h | |
| PDH E1 alpha | dehydrogenase E1 alpha subunit | Generation of Acetyl-CoA | TGGT1_245670 | 0.15 | PF3D7_1124500 | Non-mutable | no real peak | (++) 6.5 | PY00819 | (+); (+), 40 h (mid liver stage) | |
| PDH E1 beta | dehydrogenase E1 beta subunit | Generation of Acetyl-CoA | TGGT1_272290 | -0.18 | PF3D7_1446400 | Mutable | Schizont (40 h) | (++) 6.5X | PY07062 | (+);(+); 40 h | |
| PDH E2 | dehydrogenase E2 subunit | Generation of Acetyl-CoA | TGGT1_206610 | -2.97 | PF3D7_1020800 | Non-mutable | no real peak | (++) 6.5X | PY04573 | (+); (+); 40 h (mid liver stage) | |
| PDH E3 | dehydrogenase E3 subunit | Generation of Acetyl-CoA | TGGT1_305980 | -1.38 | PF3D7_0815900 | Mutable | no real peak | (++) 9X | PY00573 | (+);(+); 40 h | |
| ACCase | acetyl-CoA carboxylase | Generation of Malonyl-CoA | TGGT1_221320 | -3.6 | PF3D7_1469600 | Mutable | no real peak | (++) 23X | PY01695 | (+);(+); 50 h | |
| ACP | Acyl Carrier protein | FASII | TGGT1_264080 * | -1.43 | PF3D7_0208500 | Non-mutable | Late trophozoite-Shizont (32-40 h) | (++) 12X | PY04779 | (+); (+); 40 h | * Essential (Ramakrishnan et al. 2012) |
| FabZ | hydroxyacyl-ACP dehydratase | FASII | TGGT1_321570 | 0.67 | PF3D7_1323000 | Mutable | Trophozoite-Schizont (26-46 h) | (++) 9X | PY01586 | (+); (+); 40 h | |
| FabB/F | keto acyl-ACP synthase I/II | FASII | TGGT1_293590 | -1.21 | PF3D7_0626300 | Mutable | Ring (0 h)-Schizont (40-48 h) | (++) 9X | PY04452 | (+);(+); 40 h | |
| FabD | Malonyl-CoA transacylase | FASII | TGGT1_225990 | -0.98 | PF3D7_1312000 | Non-mutable | Ring (0 h)-Schizont (40-48 h) | (++) 7X | PY05492 | (+); (+); 40 h (mid liver stage) | |
| FabH | Ketoacyl ACP synthase III | FASII | TGGT1_231890 | -1.91 | PF3D7_0211400 | Mutable | Late trophozoite-Shizont (32-48 h) | (++) 12X | not found in *P. Yoelii* | N/A | |
| FabG | Ketoacyl ACP reductase | FASII | TGGT1_217740 | -1.89 | PF3D7_0922900 | Non-mutable | Late trophozoite-Shizont (32-48 h) | (++) 14X | PY02416 | (+);(+); 40 h | |
| FabI | Enoyl-ACP reductase | FASII | TGGT1_251930 | -1.32 | PF3D7_0615100*, ** | Non-mutable, | no real peak | (++) 11X | PY03846 | (+);(+); 40 h | * Dispensible in red blood stage (Vaughan et al. 2009) ** essential during starvation (this work) |
| LipA | Lipoic acid synthase | Lipoic acid production | TGGT1_226400 | -0.97 | PF3D7_1344600 | Non-mutable | Late trophozoite-Schizont (32-48 h) | (++) 16X | PY06208 | (+);(+); 50 h | |
| LipB | octanoyl-ACP:protein transferase | Lipoic acid production | TGGT1_315640 | -1.74 | PF3D7_0823600 | Mutable | no real peak | (++) 9X | not found in *P. Yoelii* | N/A | |
| GpdA | Glycerol-3-phosphate dehydrogenase | Glycerol-3-phosphate production | TGGT1_307570 | -3.03 | PF3D7_1216200 | Non-mutable | Trophozoite (24 h), Late schizont (48 h) | (-) 2X | PY005585 | (+); (=); 50 h (late liver stage) | |
| ATS1 | glycerol-3-phosphate acyltransferase | Phospholipid synthesis/ (L)PA production | TGGT1_270910 * | -2.13 | PF3D7_1318200 ** | Mutable | Late Trophozoite-Schizont (32-48 h) | (++) 25X | not found in *P. Yoelii* | N/A | * Essential (Amiar et al. 2016) ** Dispensible (Shears et al. 2017) |
| ATS2 | acylglycerol-3-phosphate acyltransferase | Phospholipid synthesis/ PA production | TGME49_297640 | -0.02 | PF3D7_0914200 | Non-mutable | Schizont (40 h) Rings | (+) 3X | PY02486 | N/A | |
| GPAT (ER) | glycerol-3-phosphate acyltransferase | Phospholipid synthesis/ (L)PA production | TGGT1_256980 | -2.36 | PF3D7_1212500 | Non-mutable | Late Trophozoite (32 h) | not available | PY06015 | N/A | |
| AGPAT (ER) | acylglycerol-3-phosphate acyltransferase | Phospholipid synthesis/ PA production | TGGT1_240860 * | -2.54 | PF3D7_1444300 | Non-mutable | Late Trophozoite-Schizont (32-48 h) | (++) | PY01678 | (+); (+); 50 h (late liver stage) | * Likely essential (this work) |
